# Counterfactual Hypothesis Testing of Tumor Microenvironment Scenarios Through Semantic Image Synthesis

**DOI:** 10.1101/2020.10.27.358101

**Authors:** Daniel Li, Jennifer Chen, Qiang Ma, Yubin Xie, Andrew Liu, Justin Cheung, Herman Gudjonson, Tal Nawy, Dana Pe’er, Itsik Pe’er

**Affiliations:** Columbia University; Memorial Sloan Kettering Cancer Center; Google

**Author notes:** Equal Contribution (Daniel Li, Jennifer Chen).

**Keywords:** Semantic Image Synthesis, Generative Adversarial Networks, Multiplexed Imaging, Tumor Immunology

## Abstract

Recent multiplexed protein imaging technologies characterize cells, their spatial organization, and interactions within microenvironments at an unprecedented resolution. Although observational data can reveal spatial associations, it does not allow users to infer salient biological relationships and cellular interactions. To address this challenge, we develop a generative model that allows users to test hypotheses about the effect of cell-cell interactions on protein expression through in silico perturbation. Our Cell-Cell Interaction GAN (CCIGAN) model employs a generative adversarial network (GAN) architecture to generate high fidelity synthetic multiplexed images from semantic cell segmentations. Our approach is unique in that it learns relationships between all imaging channels simultaneously and yields biological insights from multiple imaging technologies in silico, capturing known tumor-immune cell interactions missed by other state-of-the-art GAN models.

## 1 Introduction

The spatial patterns of cells in a neighborhood dictates signaling relationships and interactions with the cellular environment that are critical to development and disease. Tumor-immune cell interactions within the tumor microenvironment are a prime example, implicated in many facets of cancer pathogenesis and treatment (1; 2).

Recent advances in multiplexed tissue imaging, from fluorescence microscopy (e.g. CODEX (3), t-CyCIF (4)) to ion mass spectrometry (e.g. MIBI-TOF (5)), enable the acquisition of subcellular localizations at a high dimensional resolution (6; 7). While these emerging technologies improve resolution, methods to effectively interpret both spatial and expression-level changes in different cellular environments are lacking. The time and scale constraints of analyzing cellular interactions with incidence-based in vivo methods suggest the need for in silico methods’ speed and scale to complement. However, multiplexed imaging poses unique challenges for in silico methods due to its high-dimensionality and complex non-uniform protein signals across many channels, which, unlike typical red green blue images, do not express similar information in all channels.

Existing methods for analyzing imaging data can be broadly grouped into cell classification and population analysis, neither of which take advantage of multiplexed imaging to investigate protein localization. Methods for cell classification (8) focus on clustering cell-types by protein characteristics but do not predict protein expression and localization. Bayesian methods for cell-cell interaction analysis (9) only predict scalar magnitudes of protein expressions and lack spatial information. Traditional binning and regression-type methods can quantify neighboring cell contributions, but are limited by issues such as bin size selection, high dimensionality, feature extraction and nonlinear effects, motivating the use of alternative neural-network-based methods. Some recent GAN-based methods augment cell imaging data for training (10; 11), or conditionally model cell marker data on cell types (12; 13), but none model spatial context in multiplexed data and are not purposed to study cell cell interactions.

We develop a method that allows detailed insight into important mechanistic questions and addresses both challenges of imaging complexity and scale. To analyze subcellular protein localization and test hypotheses of cellular interactions in different microenvironments, we present Cell-Cell Interaction Generative Adversarial Network (CCIGAN). CCIGAN is a two-part framework with a novel GAN model which learns complex marker relationships and generates high fidelity multiplexed images, and a biological discovery toolkit to test counterfactual cell interaction scenarios by interpreting spatial, directional, and magnitude changes at the protein-level. CCIGAN can conditionally capture realistic but potentially manually unobserved cell scenarios. With the experimental toolkit, a user can answer mechanistic questions, such as how the expressions of inhibitory immune checkpoint receptors on a T cell change in a particular tumor environment, and search for specific cellular interactions that result in significant protein expression changes on a per-patient level.

CCIGAN successfully recapitulates well-studied patterns in tumor immuno-biology not captured by other state of the art GAN models and yields novel biological insights in immune cell-tumor interactions that are not plausible to deduce using solely ‘wet lab’ experimental perturbations. Our model generalizes across multiple high-dimensional multiplexed technologies and cancer types and provides insights into cellular interactions in disease pathogenesis that cannot be achieved by direct observation of data alone.

## 2 Results

CCIGAN generates high fidelity multiplexed cell images using a novel conditional protein mechanism that modulates protein expressions based on cell types, morphology, and neighborhood. We find CCIGAN rediscovers cell-cell interaction phenomena supported by previous wet-lab studies while previous state-of-the-art GANs learn incorrect trends. CCIGAN’s ability to recapitulate biological phenomena also generalizes across multiple high dimensional imaging technologies (MIBI-TIF, CODEX, t-CyCIF), cancer types, and patient characteristics. Finally, leveraging CCIGAN, our search algorithm discovers highly significant subcellular mechanisms and allows for hypothesis testing of potentially new cellular phenomena.

### The CCIGAN Model

As a deep conditional image synthesis model, CCIGAN takes as input labeled cell segmentations patches (subsection of a full deep-learning based cell segmentation usually included in the dataset) and generates a multiplexed cell image where each channel is a spatial prediction of a particular protein being expressed on the cells in the segmentation patch (Figure 1; see Online Methods 5.2 for data format and processing).

**Figure 1:**
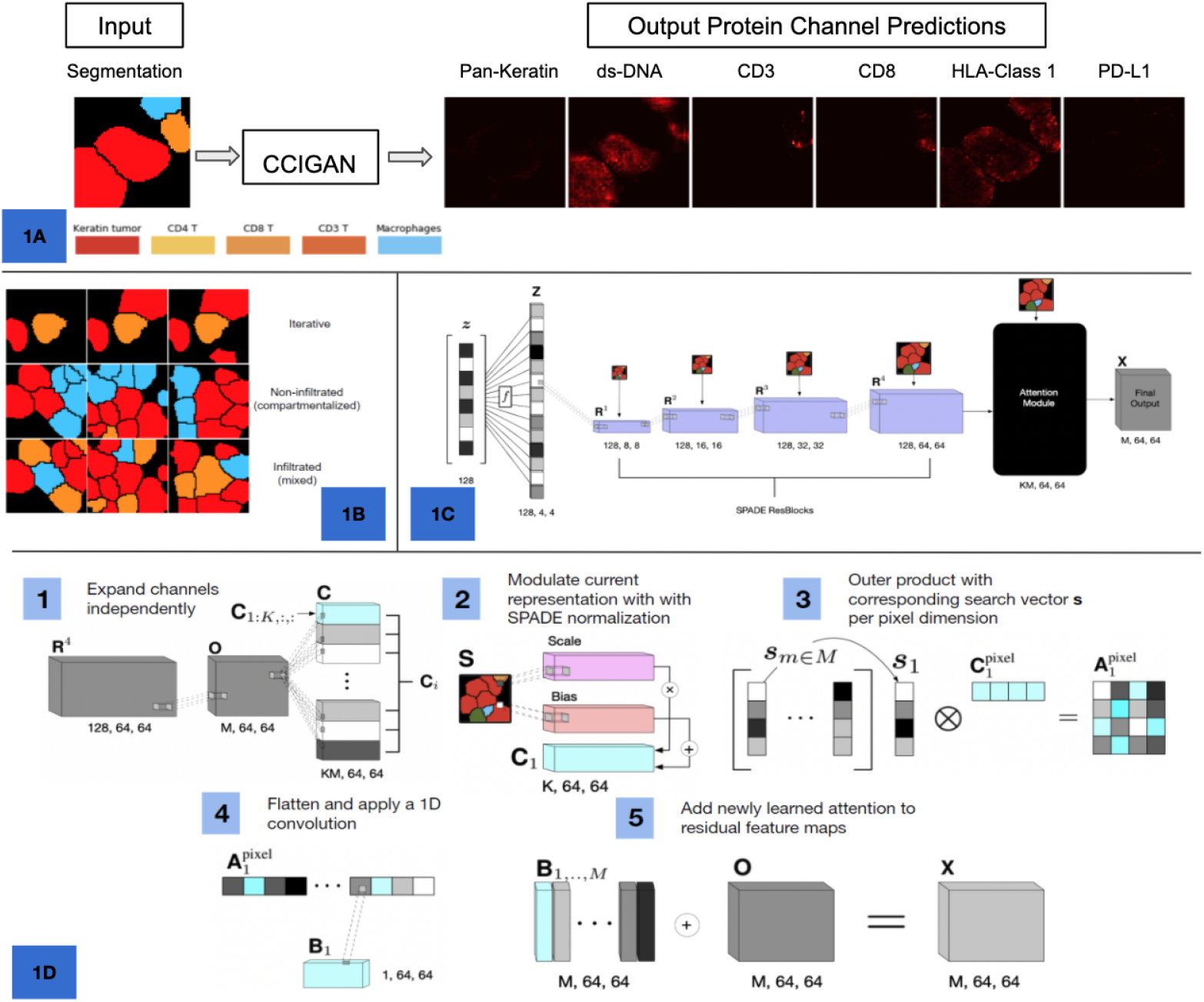
CCIGAN model overview and components. **(A)** CCIGAN takes a labeled segmentation patch as input and returns predicted expression values (between 0 and 1) in each channel. **(B)** CCIGAN segmentation maps can be synthetically altered to pose different counterfactual biological scenarios for in silico hypothesis testing. Example scenario in the first row tests the effect on a CD8 T cell (orange) of adding tumor cells (red) around it. Second and third row pose different immune infiltration scenarios. **(C)** CCIGAN convolutional architecture (see Online Methods 5.1.1 for equations). A low dimensional noise vector (z) is convolutionally upsampled and normalized by an input segmentation through a series of spatially adaptive normalizing (SPADE) res-net blocks. The intermediate representation passes through a multiplexed attention module to reason biological relationships. **(D)** Attention module description (see Online Methods 5.1.2 for equations). The module first disentangles the fully connected representation into individual channels through grouped convolutions, then uses an outer product attention layer to model relationships and interactions between specific markers and cell types. Using an outer product forces attention at a pairwise pixel-level comparison for all combinations of a learned prior vector over markers and different cell types.

CCIGAN learns a many-to-many mapping between different cell types and different protein markers with a novel protein attention mechanism. Due to multiplexed data’s unique challenge of non-uniform, relatively independent biological channels, generating multiplexed imaging is difficult. Conventional GANs for RGB images equate protein channels to be identical distributed across different cell types and place equal generative weight on every pixel in the segmented cells. CCIGAN overcomes this challenge by independently conditioning each generated protein channel on the cell types in the input segmentation (Fig. 1D) and up or down-weighting channel expressions based on cells’ types, shape, or relative locations in the segmentation. For example, for the protein PD-1 (typically expressed only in immune cells), the attention module focuses on generating the subcellular pattern of the immune cell’s PD-1 while ignoring irrelevant cells.

CCIGAN is unique among biological tools because current cell image analysis methods do not perform multichannel or multicellular image synthesis or are limited in the number of proteins and complexity of cell-cell interactions it can model. CCIGAN’s architecture is novel in conditioning spatial protein expression on a variety of cell types, morphology, and relative locations. It accomplishes this by building on top of convolutional networks with spatially adaptive normalization (SPADE) (14) and a novel protein attention mechanism (Fig. 1C, Online Methods 5.1.1). SPADE conditions protein expression distributions per cell types in the input patch, allowing the model to use many types of cell information to predict spatial protein expression. Our attention mechanism then further refines spatial expression importance by learning important protein expression localizations to up or down-weight. This is accomplished through persistent protein vectors that interact with SPADE’s outputted conditional feature maps using outer products, convolutions, and transformations (S1.1). A technical exploration into model interpretability of the protein attention vectors is also given in S1.4. Due to its conditional nature, CCIGAN is also able to take user-defined segmentations with deliberately manipulated cell types and shapes as input, allowing for robust hypothesis testing and experimental analysis. CCIGAN works best with high dimensional multiplexed imaging (500 nm/pixel resolution or better) because its deep convolutions require granular input signals.

### Model evaluation

We evaluate CCIGAN against other state-of-the-art image synthesis methods on the multiplexed ion beam imaging (MIBI) triple negative breast cancer dataset (6) and find it generates images that are more biologically sound. 15 patients in the MIBI dataset with greater than 10% frequency of CD8 T cell and tumor cells, and more than 4500 cells/patches in total (top 40% percentile) were evaluated in two ways. First, using standard computer vision metrics to evaluate image reconstruction of protein channels, we report that CCIGAN outperforms in 4 of 6 reconstruction metrics and is matched by SPADE in L1 loss (Table 1, Online Methods 5.3.1). Most notably, CCIGAN outperforms in Cell Mutual Information (S3.3) between the predicted and ground truth cells, indicating it succeeds in generating realistic protein distributions. Second, we benchmark models’ ability to synthesize biological phenomena using our biological discovery toolkit (Fig. 2). CCIGAN is able to recapitulate established patterns of cell-cell interactions linked to known biological phenomena while other models fail to learn correct cell-cell interaction associations (Fig. 3).

**Table 1:** Comparison of conventional reconstruction metrics between different models. Higher SSIM and Mutual Information, and lower L1 and MSE indicate better model.

| Metrics | CCIGAN | SPADE | Pix2PixHD | CycleGAN |
| --- | --- | --- | --- | --- |
| Adjusted $L_1$ Score | **1.199** | **1.085** | 1.484 | 5.656 |
| $L_1$ Score | **1.137** | **1.067** | 1.326 | 4.744 |
| Adjusted MSE Score | **0.067** | **0.069** | 0.159 | 2.286 |
| MSE Score | **0.066** | **0.068** | 0.153 | 1.907 |
| SSIM | **0.799** | **0.786** | 0.676 | 0.429 |
| Cell Mutual Information | **10.165** | 9.172 | 8.068 | 6.387 |

**Figure 2:**
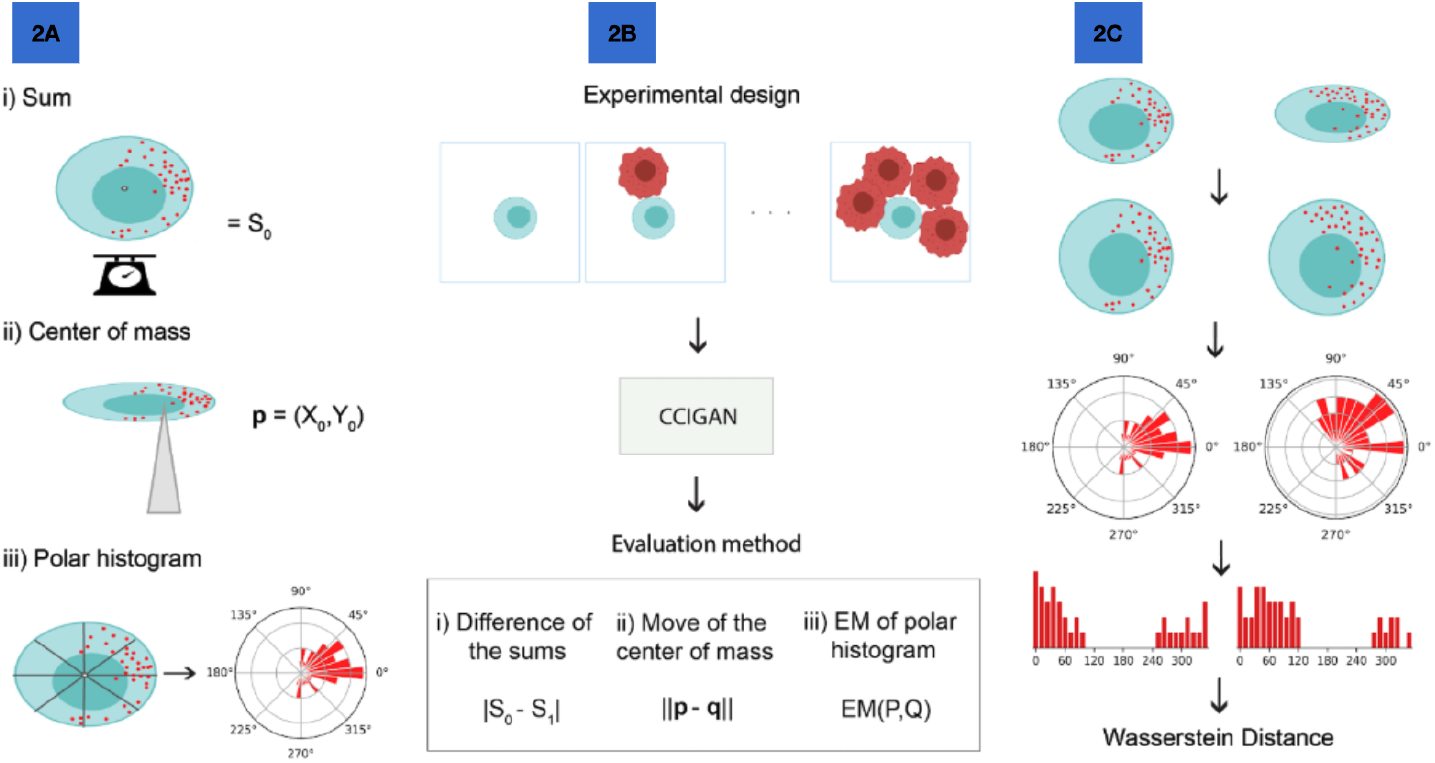
Biological Discovery Toolkit. **(A)** We designed metrics to evaluate changes in protein expression (red dots) within a cell of interest (COI) (blue circle) due to changes in cell neighbors: (i) total protein expression in cell of interest, (ii) center of mass of a protein expression within cell, and (iii) spatial shift in expression. **(B)** To explore unknown interactions, we provide an automated **search algorithm** that scans CCIGAN predictions for a wide range of cell types and segmentation arrangements, and assesses changes in a specified metric. Red circles indicate a different cell type. **(C) EM (Earth Mover’s) Score**. To evaluate differences in spatial-directional protein expression (directional mass movement), the cell is first warped into a standardized format with interpolated expression values, values are then binned, and a histogram is computed along its polar axis; finally, center of mass geometries are used as guiding directions to positively or negatively weight the earth mover’s distance in computing the EM Score. Positive EM Score indicates a cell’s protein shift towards another cell whereas a negative EM Score indicates a protein shift in the opposite direction.

**Figure 3:**
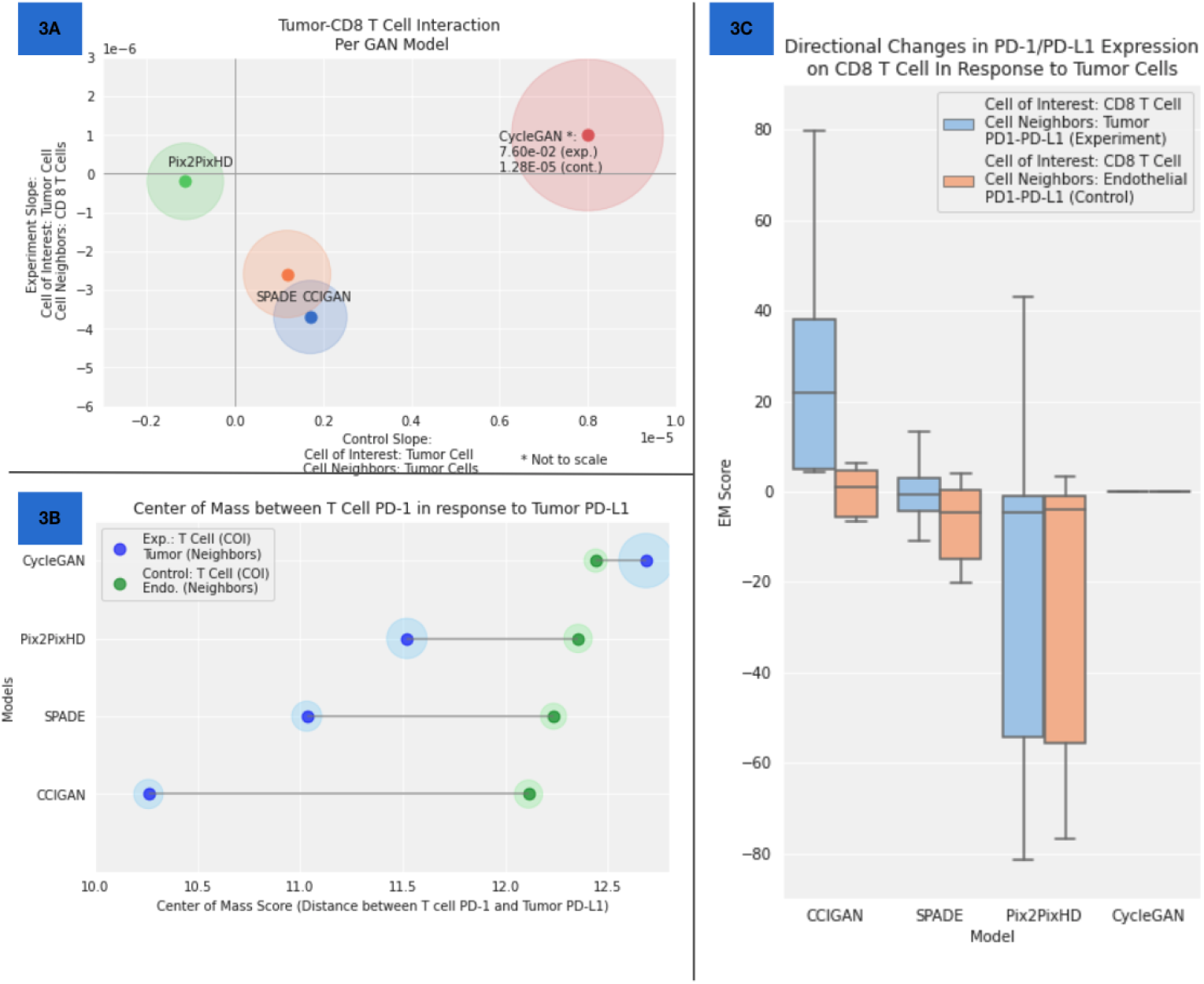
Evaluation of CCIGAN compared to existing GANs. Using three main experiments from our biological discovery toolkit, we test inference of cell-cell interactions from MIBI images of triple negative breast tumors across 15 patients. **(A)** Predicted effects of neighboring CD8 T or endothelial control cells on pan-keratin levels in a tumor cell of interest. When CD8 T cell’s surround a tumor cell, CCIGAN and SPADE both accurately predict a decrease in total pan-keratin expression, but CCIGAN predicts a more negative trend with lower variance across patients. In a control where a tumor cell is surrounded by tumor cells, all models other than CycleGAN accurately predicts no significant trend in pan keratin. Slopes represent rates of change in pan-keratin as a function of the number of cell neighbors. 50% confidence intervals (CIs) between patients are reported in lighter background circles. **(B)** Differences in predicted center of mass (COM) between T-cell-expressed PD-1 and tumor-cell-expressed PD-L1. Larger differences between the model (blue) and control (green) scores indicate the model is better at capturing the true response of PD-1 in a real scenario. CCIGAN reports the smallest COM score for the experimental scenario, indicating that T Cell PD-1 is expressed close to Tumor PD-L1, and it reports the largest difference between experimental and control. 95% CIs are reported in background circles. **(C)** Differences in predicted spatial shift of protein expression. Higher Earth Mover’s (EM) Scores (15) reflect stronger shifts. Predicted PD-1 expression shift in a T cell of interest resulting from PD-L1-expressing tumor cell neighbors or control endothelial cells. CCIGAN supports the tumor PD-L1 and CD8 T cell PD-1 interaction with large positive EM Score for the experimental scenario, whereas other models capture small to negative EM scores.

### Biological Discovery Toolkit

In order to evaluate the biological utility of CCIGAN and answer questions about cellular mechanisms in the data, we develop a series of biological interpretability techniques for estimating cell-cell interaction hypotheses and a search algorithm for discovering significant cell-cell interactions.

The first part of our biological discovery toolkit comprises of techniques for counterfactual hypothesis testing of cell-cell interactions (Table 2). These techniques quantify how one cell’s protein expression at a pixel level reacts to newly introduced adjacent cells (Fig. 2). We can ask questions such as “how would a tumor cell be affected by adding CD8 T cells next to it?” (Fig. 1B) by artificially inserting adjacent T cells in the segmentation patch and observing the change in predicted protein expression on the tumor cell. CCIGAN’s conditional and generative nature allows for hypothesis testing of user-manipulated cell patches with different cell types, location and morphology. CCIGAN can discover significantly correlative cell-cell interactions in a dataset but is not a replacement for in vivo or wet lab testing to establish causality. However, such hypothesis testing is still important as manipulating original data to investigate protein expression changes is not sufficient or efficient. Deleting or isolating cells would not capture changes in protein expressions due to the removed cell (see S2.4 for an illustration). We assess our toolkit on the same 15 MIBI patients as the previous section and evaluate all generative experiments (not requiring comparison with the original protein distributions) on a test set of 1000 segmentation patches that are manipulated into hypothetical scenarios by changing cell types in original test set segmentation patches.

**Table 2:**
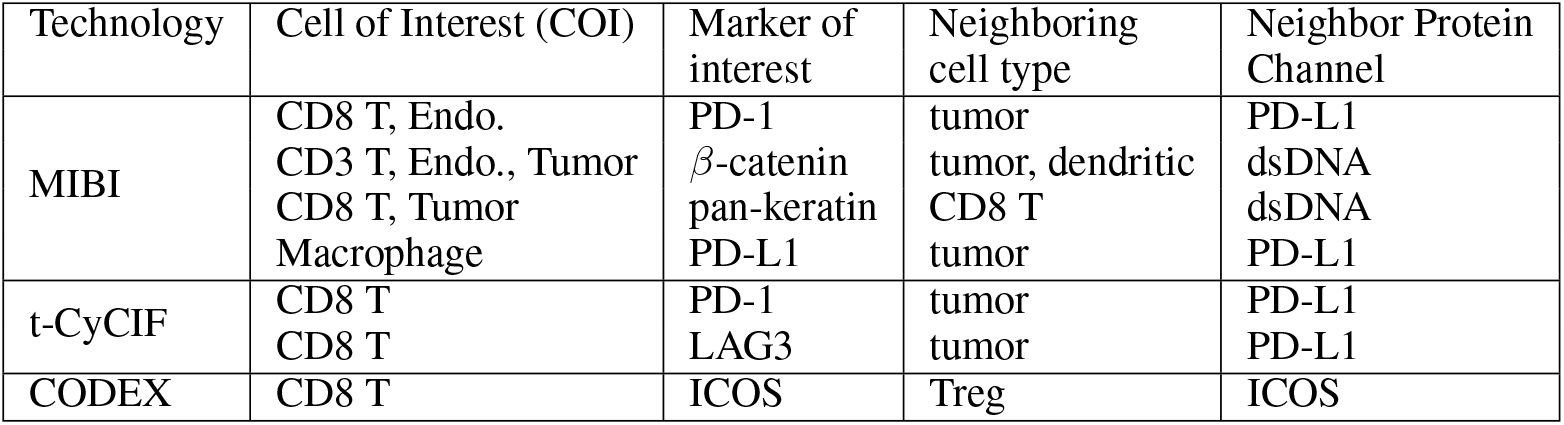
Rediscovery of known cell-cell interactions as positive controls for CCIGAN

First, we examine expression trends in the cell of interest (COI) as we iteratively add neighboring cells. In the MIBI dataset (6) we examine trends in pan-keratin levels when CD8 T cells mediate tumor killing. Previous literature states that CD8 T cells release granzymes or induce extrinsic apoptosis which results in the release of enzymes that cleave the tumor cell’s pan-keratin (15), disrupting cell structure (16; 17; 18; 19). We explored how a drop in pan-keratin levels in a tumor cell of interest could be used as a proxy for the likelihood of tumor cell death (prior to observed cell destruction) in the presence of adjacent CD8 T cells (Fig. 3A, Online Methods 5.3.4, S3.4). CCIGAN and SPADE both predicted decreases in tumor pan-keratin expression that became more dramatic when surrounded by more CD8 T cells, whereas introducing adjacent tumor cells (control) produced little change. CCIGAN predicted a more negative trend than SPADE, while other methods failed to detect a negative relationship. CycleGAN (20) erroneously reported very large changes under experimental and control conditions.

Next, we examined a cell’s center of mass (COM) shift in response to adjacent neighbors. In MIBI, we assess PD-1 expression in a CD8 T cell of interest is affected by adjacent tumor cells (Fig. 2A.ii, Fig. 3B, Online Methods 5.3.2, S3.1). T cells located within the tumor microenvironment upregulate PD-1 expression as a result of influences from the tumor milieu (21; 22; 23). The T cell’s PD-1 center of mass (COM) was used as a proxy for its localization, and adjacent tumor cells’ PD-L1 center of mass was used as a proxy for cell contact. The score represents the distance between the PD-1 and PD-L1 centers of mass in pixels, with greater predictive accuracy represented by lower scores, since shorter distances are expected for this direct protein-protein interaction (13). Larger differences in score between model and a randomized control indicate the model is better at capturing the true PD-1 response. Using these metrics, CCIGAN predicts better localized PD-L1 expression than other methods such as SPADE (14) and Pix2PixHD (24), while CycleGAN predicts the opposite biological insight. CCIGAN also outperforms SPADE in predicting a lower distance between T cell PD-1 and Tumor PD-L1 COMs.

Third, we test CCIGAN’s ability to predict directional mass shift (as Earth Mover’s or EM) of protein localization in response to neighboring cells. We hypothesize that iteratively introducing adjacent PD-L1-expressing tumor cells will induce directional shifts in PD-1 expression in a T cell of interest (Fig. 2A.iii, Fig. 3C, Online Methods 5.3.3). An Earth Mover’s (EM) Score with a higher magnitude indicates a stronger cell-cell interaction; positive scores indicate the direction of protein localization moving towards cell neighbors and negative values otherwise. CCIGAN confirmed biological expectation by generating a large positive EM Score, indicating a shift in T-cell PD-1 expression towards the PD-L1 center of mass in adjacent tumor cells (Fig. 3C). In contrast, the addition of control endothelial cells only resulted in a small shift in T-cell PD-1 expression. PD-L1 is found on a variety of endothelial cell types and serves as an important immune checkpoint that protects normal cells from T-cell-driven autoimmune reactions (25; 26); thus, CCIGAN results for the endothelial control also conform to biological expectation. The difference in magnitude and directional effect of T-cell interaction with tumor and endothelial cells highlights the greater degree to which the PD-1/PD-L1 immune checkpoint is exploited by malignant cells to escape immune detection. Off-the-shelf methods SPADE, Pix2PixHD, and CycleGAN fail to capture this interaction and often predict negative experimental EM shifts.

The second component of our biological discovery toolkit is our search algorithm, which identifies significant cell-cell interactions (Online Methods 5.4) by repeatedly testing different counterfactual cell-cell interaction scenarios and evaluating how frequent the scenario induces changes in protein expression. On MIBI data for a single patient, the search algorithm finds many well-established cellular mechanisms (e.g. PD-1/PD-L1 in response to CD8 T cell of interest and Tumor neighbors) (Online Methods Fig. 5). However, there are some erroneous correlations (e.g. a CD8 T cell expressing PD-L1 while in close contact to tumor cells) that are possibly attributed to poor segmentations, imaging faults, and other noisy effects. These limitations underscore the use of CCIGAN as a complement to traditional experimental methods to enable faster hypothesis testing and discover overlooked cellular mechanisms that warrant further investigation. A further discussion into significant cell-cell interactions discovered by the algorithm, (in particular vimentin-related interactions), is presented in S4.2.

### Generalizability across imaging technologies

We present additional analyses on three imaging technologies to corroborate previously established biology and demonstrate CCIGAN’s generalizability across many technologies, diseases, and patient characteristics.

### Applications to MIBI data

On MIBI data, we applied an analysis using EM (similar to Fig. 3C) on *β*-catenin expression directional change in dendritic cells (DCs) upon interacting with tumors. Wnt signaling from tumors have been found to induce *β*-catenin expression in dendritic cells (DCs), suppressing DC activation (28). Increasing tumor cell presence adjacent to DCs triggers downstream immunosuppressive effects of the Wnt pathway, increasing directional *β*-catenin expression in the DC cell. To quantify changes of DC cell *β*-catenin expression due to neighboring cells, we measure mass movement between dsDNA expression of neighbors (indicating magnitude of cell presence) and a DC cell’s *β*-catenin expression (Fig. 4A). Using CCIGAN trained on one patient, we add tumor cells next to a DC cell of interest resulted in a directional increase in *β*-catenin (EM Score 333.9) that is greater than that observed upon adding endothelial cells (EM Score 233.6) or CD3 T cells (EM Score 89.3). These findings confirm that CCIGAN detects expected directional patterns of the Wnt pathway activation in DC cells.

**Figure 4:**
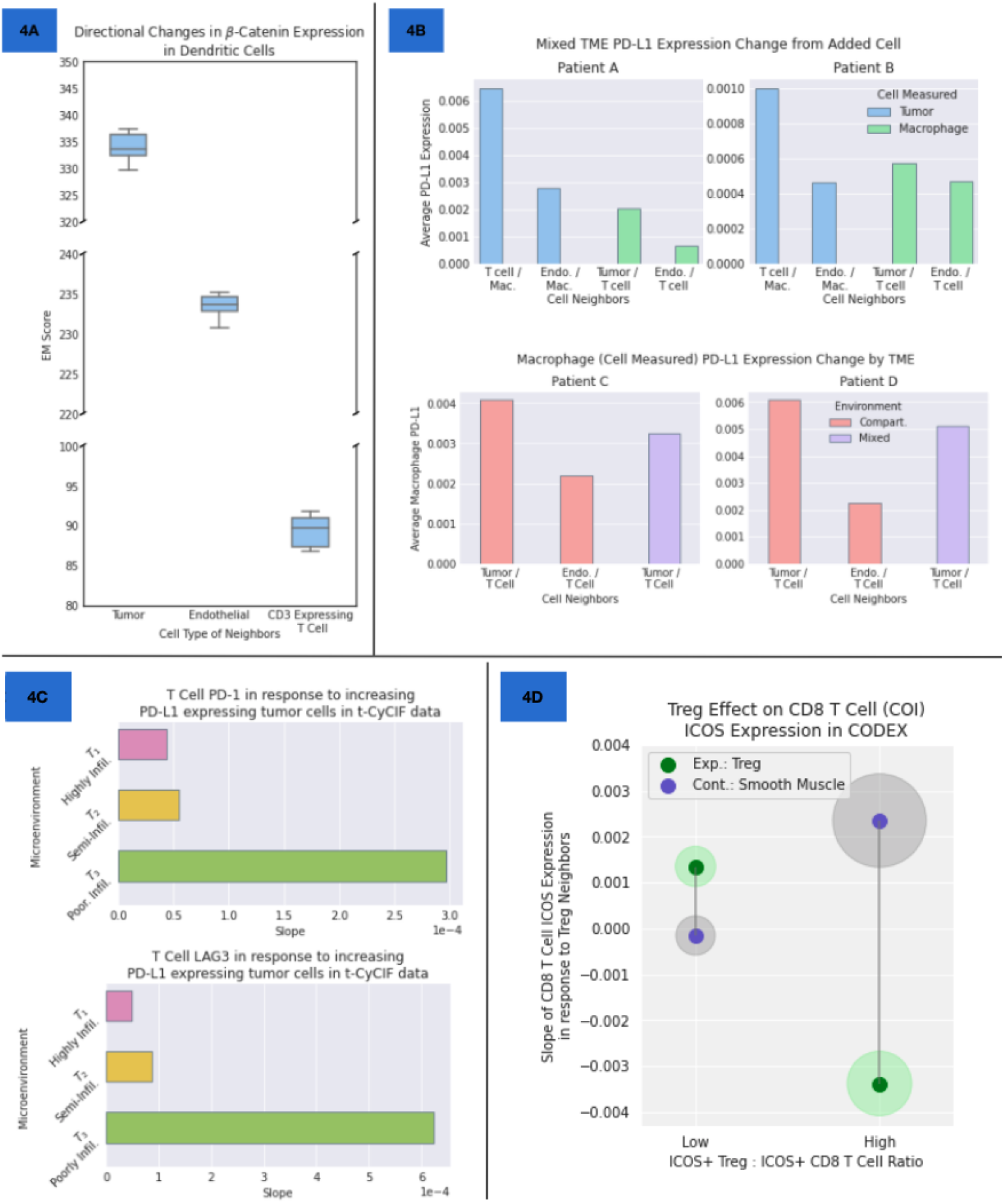
Recapitulating discovered cell-cell interactions. **(A)** Predicted shifts in directional *β*-catenin expression in dendritic cells (DCs) on adjacent tumor vs non-tumor cells. CCIGAN predicts high EM Score of DC cell *β*-catenin expression with a tumor cell neighbor, suggesting strong Wnt activation in the DC cell and confirming biological findings on the tumor’s immunosuppressive effects on the DC. **(B)** Effects of tumor compartmentalization in tumor-immune cell interactions (1B Row 2), evaluated on MIBI data from TNBC patients in Keren et al. 200 test neighborhood segmentations (90-10 train/test split) were generated to simulate different environments. Control/reference: endothelial (endo.) cell. **(Upper)** Within the mixed tumor microenvironments experimental set up, PD-L1 expression in tumor and macrophage cells is induced by T cells & macrophages or tumors & T cells respectively, representing infiltrated neighborhoods. The PD-L1 expression in tumor or macrophage cells (legend) was measured next to their appropriate cell neighbor combinations (*x*-axis). **(Lower)** Macrophage PD-L1 expression increases induced by neighboring tumor cells within compartmentalized and mixed cell environments. Greater PD-L1 expression increase in macrophages in the compartmentalized environment than mixed environment recapitulates previous literature. **(C)** CCIGAN inference of cell-cell interactions from t-CyCIF images of primary lung cancer (27). **(Upper)** PD-1 expression in CD8 T cells as a function of number of PD-L1 expressing tumor cell neighbors and type of tumor environment. Slope is calculated similarly to 3A. In an infiltrated environment (high lymphocyte presence within the tumor microenvironment, indicating strong anti-tumor immune response), reduced PD-1 expression on infiltrating T cells signals a more robust anti-tumor immune response. **(Lower)** LAG3 expression in CD8 T cells as a function of PD-L1 expressing tumor cell neighbors and type of tumor environment. In an infiltrated environment, a decrease in T cell LAG3 expression indicates reduced T cell exhaustion in situations where there is a robust anti-tumor immune response. **(D)** Iterative trend experiment to investigate Treg’s immunosuppressive effects on CD8 T cell ICOS expression in CODEX colorectal cancer data. In patients with low ICOS+Treg to ICOS+CD8 T cell ratio, additional Treg neighbors do not downregulate CD8 T cell ICOS expression, whereas in patients with high Treg/CD8 ratio, more Treg neighbors is associated with decreased T cell ICOS expression. 95% CIs across patients are reported in background circles.

**Figure 5:**
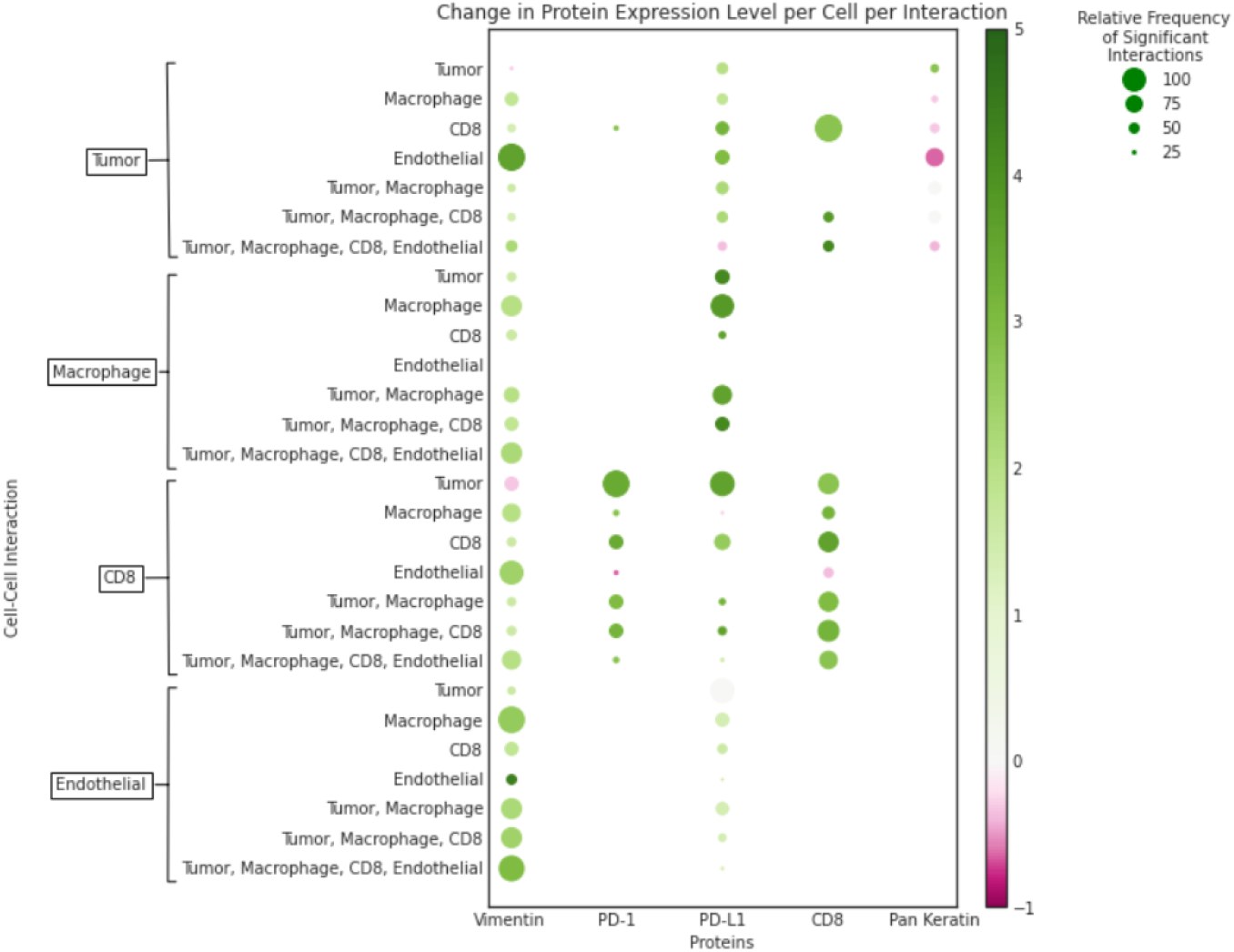
Search algorithm results. Specific cell-cell interactions and the associated magnitude of change in protein expression. The cell types in boxes on the y-axis represent the cell of interest in each experiment; cell names adjacent to these boxes represent neighbors. The circle diameter represents the relative frequency of this specific cell -cell interaction inducing a non-trivial change in the expression of a given protein. The hue of each circle denotes the order magnitude of change in expression level. For example, a CD8 T Cell (boxed, y-axis) with tumor cell neighbors (y-axis) experiences frequent and significant interactions that induce a change in the T cell’s PD-1 expression (x-axis). These interactions are predicted to induce an average of 3 orders of magnitude increase in PD-1 expression.

In another study across TNBC patient cohorts (Fig. 4B, Online Methods 5.3.5, S3.5), CCIGAN corroborated Keren et al.‘s predictions on the PD-1/PD-L1 pathway and the variability of PD-L1 expression in macrophage and tumor cells in different TMEs. In compartmentalized TMEs, tumor and immune cells are spatially segregated, whereas in mixed TMEs, tumor and immune cells are mixed together. We trained 4 individual models on 2 exemplar patients chosen per TME type^1^ from the TNBC cohort to demonstrate CCIGAN’s ability to learn cell-cell interactions dependent on specific TMEs. In the mixed environment (Fig. 4B Upper), CCIGAN predicted higher increases in PD-L1 expression in tumors and macrophages in T cell infiltrated neighborhoods than other control neighborhoods containing endothelial cells. Reiterating Keren et al.‘s finding in mixed TMEs, we also observe tumors expressing the highest average PD-L1 expression, substantially higher than macrophage PD-L1 expression, in their infiltrated neighborhoods. We further corroborate Keren et al.‘s second claim of macrophages exhibiting the highest PD-L1 expression in compartmentalized TMEs (Fig. 4B Lower). Models trained on compartmentalized patient data were used to evaluate macrophage PD-L1 expression on both compartmentalized and mixed neighborhoods and predicted greater PD-L1 expression in macrophages that reside in tumor compartments as opposed to mixed environments (Fig. 4B Lower). This phenomenon further reveals CCIGAN’s capability to learn general biological interactions even if trained on different TMEs.

### Applications to Tissue Cyclic Immunofluorescence (t-CyCIF)

t-CyCIF is a novel iterative immunofluorescence staining procedure that produces high resolution imaging (4). We apply CCIGAN to a t-CyCIF primary lung cancer image (27) that exhibited progressive tumor cell growth. To test CCIGAN’s ability to learn cell-cell interactions present in different TMEs, we split the image data according to three different stages of tumor-infiltration by immune cells – poorly-infiltrated (*T*_1_), semi-infiltrated (*T*_2_), and heavily-infiltrated (*T*_3_) (S2.2.2) and trained one CCIGAN model per microenvironment.

For each of the three microenvironment-specific models, we examined the correlation between T cell (COI) PD-1 and tumor PD-L1 (Fig. 4C Left, S3.6) using the slope trend experiment (Fig. 3A). In the non-infiltrated microenvironment model, we observe a substantially higher trend of T-cell PD-1 expression as a function of tumor area than in the semi-infiltrated which, in turn, was also slightly higher than the heavily-infiltrated microenvironments. These findings are consistent with the idea that tumors with poor lymphocyte-infiltration have had greater success in suppressing anti-tumor immune responses and thus, an increase in PD-1 would be expected in the T cell (cell of interest).

Similarly (Fig. 4C Right, S3.6), we examined the association between exhaustion marker LAG3 (29; 30) expression on T cells, and PD-L1 expression on neighboring tumor cells. CCIGAN predicted an increase in LAG3 expression on the T cell (cell of interest) as tumor neighbors were added, with greater levels of induction from infiltrated to semi-infiltrated to non-infiltrated tumor scenarios. These results agree with previous studies that increased tumor PD-L1 expression suppress T cell function and induce T cell exhaustion, and that T cells within tumor environments of higher lymphocyte-infiltration would have lower LAG3 expression, indicating less T cell exhaustion.

### Applications to CODEX

Lastly, we apply CCIGAN to a CODEX colorectal cancer (CRC) dataset (3) and explore the role of regulatory T (Treg) cells in immunosuppression of CD8 T cells. For patients with severe symptoms (diffuse inflammatory infiltration, DII), Schürch et al. (3) found local enrichment of ICOS+ Treg cells, Ki-67+ CD8, and ICOS+ CD8 T cells in tumor boundary neighborhoods. They suggest that Treg cells could “oppose the cytotoxic activity from Ki-67+ and ICOS+ CD8 T cells”, therefore causing the DII cohort to have impaired survival.

We evaluate immunosuppresion in DII patients by investigating ICOS (inducible costimulator): a marker indicating T cell activation when expressed in T effector cells, but strong immunosuppression when expressed in Tregs (31). Given ICOS’s dual function depending on the cell type it is expressed in, the ratio of Tregs to CD8 T cells is an important prognostic factor in cancer and high Treg/CD8 ratio has been observed to impair patient survival (32), perhaps due to greater immunosuppression. In DII patients with high frequency of both ICOS+ Tregs and ICOS+ CD8 T cells (6 patients, greater than 20 Tregs and CD8 cells each), we train a CCIGAN model per patient and employ the iterative trend experiment (Fig. 3A) to investigate CD8 T cell (COI) ICOS expression as a function of additional Treg neighbors (or smooth muscle as controls). In patients with low ICOS+ Treg to ICOS+ CD8 ratio (0 *< r <* 0.5), CCIGAN learned a slight positive trend similar to the control slope of the CD8 T cell’s ICOS expression as a function of additional Treg cell area. However, in patients with high ICOS+ Treg/CD8 ratio (0.5 ≤*r <* 1.0), we observed a strong negative trend that is divergent from the control slope (Fig. 4D). This suggests that in patient TMEs with higher Treg/CD8 ratio, CCIGAN accurately predicts Tregs’ immunosuppresive effects on CD8 T cell and its ICOS expression, whereas in TMEs with low Treg/CD8 ratio, the immunosuppressive effects are not apparent.

## 3 Discussion

As multiplexed imaging technologies continue to rapidly increase in resolution and availability, tools to analyze and interpret their outputs must evolve. To address limitations of current methods, we introduce a novel protein attention image synthesis model, CCIGAN, and an accompanying biological discovery toolkit to interpret CCIGAN’s learned biological mechanisms. By simultaneously conditioning on cell types, location, and morphology to learn spatial protein expression, CCIGAN then allows users to pose questions about counterfactual cell-cell scenarios in a test environment and estimate their resulting protein localization and behavior.

CCIGAN’s capacity for generating subcellular protein predictions represents a step forward in modeling cellular relationships within different microenvironments. Rather than assessing in vivo incidence of cell interaction phenomena, CCIGAN allows for dynamic hypothetical biological situations to be generated and analyzed. This addresses the difficulties of scaling in vivo experiments and expands the power of individual datasets by complementing in vivo experimentation with computational hypothesis testing.

Although CCIGAN represents a promising rapid hypothesis testing tool for in silico prediction of cell-cell interactions, guided in vivo experiments are still needed to corroborate CCIGAN’s quantitative predictions. Limitations exist in modeling, such as noisy cell segmentations and cell type classifications in the input data and possible hallucination of cell-cell interactions due to the nature of generative modeling. While the model performs well in learning and associating clear trends, rare cell-cell protein interactions may also be interpreted as noise or with low confidence.

Nonetheless, we have shown that CCIGAN outperforms state-of-the-art image synthesis methods, and is able to capture, recapitulate, and quantify clinically established cell-cell interactions. Furthermore, we evaluate CCIGAN’s ability to generate well-studied biological phenomena across multiple disease contexts, patients, imaging technologies, and TME characteristics. Given CCIGAN’s strong corroboration of previous clinically established biology and the utility of giving users freedom to test new interactions to guide further experiments, we expect this work to be useful in the multiplexed imaging domain.

## 4 Code Availability

Our modeling and preprocessing code is available for download here. To download the datasets we used, please refer to their original papers (33; 27; 3).

## 5 Online Methods

### 5.1 Methods

#### 5.1.1 Model Overview

For CCIGAN, we use SPADE residual blocks (14) as our generative backbone and DCGAN’s discriminator’s architecture (34). (14) have shown SPADE to be an effective way to inject conditioning into a generative model. The SPADE normalization layer serves as a replacement for previous layer normalization techniques. Instead of learning a universally shared per channel affine transformation, like in Batch or Instance Normalization, SPADE learns to predict affine transformations based on segmentation maps; each feature is uniquely transformed based on its cell type, size, and neighboring cells. The ability for SPADE to modulate activations based on the context of adjacent cell segmentations allows the network to effectively model the behaviors and interactions of cells. The input of CCIGAN is a noise vector ***z*** ∈ℝ^128^ and a segmentation map **S**. *f* denotes a linear layer ℝ^128^ ↦ℝ^2048^. **R**^*i*^ are feature map representations from SPADE resblocks and **X** denotes the final output of *M* cell expressions. Below, each layer’s output dimensions are given next to their respective equations. Further details such as kernel size, activation functions, training regimen, and model interpretability are given in S1.1.

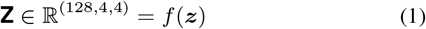

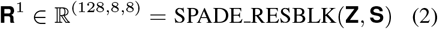

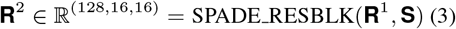

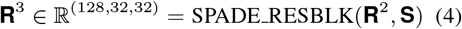

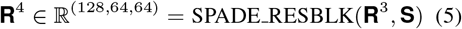

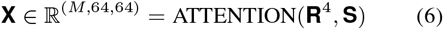

Our architectural contribution is a protein marker dependent attention module in the final output layer. The goal of the attention module is to condition the final output of a channel on a protein marker *m* and **S**’s cell types. For example the protein marker, pan-keratin *m*_pk_, is expressed exclusively in tumor cells but not in other cells. Appropriately, an attention mechanism should attend to tumor cells and ignore irrelevant cells in **S** for *m*_pk_. To replicate a marker searching for specific cell types that express it, we define a learned persistent vector for each marker denoted by ***s***_*m∈M*_ *∈*ℝ^8^ that undergo a series of operations with the final feature map representation attending to *m*’s specific cell types. It is also worthwhile to note that these persistent vectors ***s***_*m*_ offer a degree of model interpretability that mimic real world markers. The current input dimensions to the attention module are ℝ ^(128,64,64)^ following the last resblock **R**^4^ and *m* indexes from 1, .., *M*.

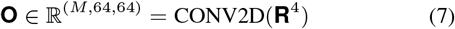

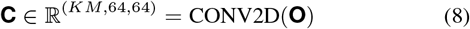

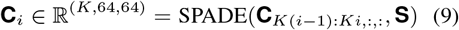

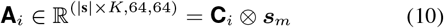

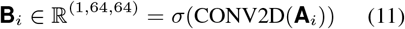

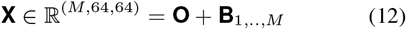

After **R**^4^, a bottleneck convolution is applied to match the original data’s dimension as **O** (step 1), which is used in a residual manner with the final output. Intuitively at this stage, **O**’s feature maps resemble the target **Y**, but we wish to further refine the output channels. We convolve **O** into *MK* channeled features for each protein marker where *K* = 8. Considering each **C**_*i*_ where *i* ∈{1, …, *M*} as a group of *K* channels, the model spatially adaptive normalizes each **C**_*i*_ and computes an outer product with the corresponding persistent vector *s*_*i*_ and **C**_*i*_. The resulting matrix is flattened and convolved (with a kernel size of 1 on the pixel level) from **A**_*i*_ *∈* ℝ ^(|**s**|×*K*,64,64)^ ↦ ℝ ^(1,64,64)^ followed by a sigmoid *σ*(·) activation. Lastly, the attentions **B**_1,…,*M*_ are added to **O** to obtain the output **X**.

Initially, the model has no priors over the interaction of protein markers and cell types. The proposed outer product attention layer (outer product and 1 × 1 convolution) excels at modeling these relationships and interactions between specific markers and cell types. By using an outer product, the model forces attention at a pairwise pixel level comparison for all combinations of elements between ***s***_*m*_ and **A**_*i*_. As training progresses, both the learned features over segmentation patches and the learned persistent vectors ***s***_*m*_ improve, in turn allowing the learned 1 × 1 convolution to reason about positive or negative relationships from the pairwise pixel combinations.

### 5.2 Data and Data Processing

We trained CCIGAN on two types of cell data, MIBI-TOF (multiplexed ion beam imaging by time-of-flight) and t-CyCIF (tissue cyclicimmunofluorescence) data. While these two types of multiplexed image data were obtained through different procedures, they share structural similarities.

Multiplexed cell images display multiple protein marker expression levels. These images are represented as high dimensional tensors **T** ∈ ℝ ^(*M,H,W*)^, *M* being the number of markers, *H* being the height of the image, and *W* being the width. Each of these markers *m ∈* {1, …, *M*}, are given as a channel taking on real values continuous in [0, 1] at each (*x, y*) coordinate, indicating the expression level at a given protein. Protein markers’ particular expression levels (either separately or in conjunction with other protein markers) demarcate different cellular subtypes and furthermore, are indicative of the functional properties of a cell. By simultaneously imaging over multiple protein markers, multiplexed images are able to identify cell type as well as provide detailed information of sub-cellular structure, cell neighbors, and interactions in the tumor microenvironment across these different marker settings.

#### 5.2.1 MIBI-TOF

MIBI-TOF images are represented in **T** ∈ℝ ^(M,2048,2048)^. These images are then further processed at a cell by cell basis into **Y** ∈ℝ ^(M,64,64)^ patches, where a cell is at the center of the patch along with its neighbors. Next, we construct semantic segmentation maps **S** ∈ ℝ ^(C+1,64,64)^, where a vector **S**_:,*i,j*_ is one-hot encoded based on a cell type *C* = 17, and the *C* + 1-th channel denotes empty segmentation space. The data is train-test split at a 9:1 ratio at the MIBI-TOF image level to avoid cell neighborhood bias.

Data obtained through MIBI-TOF characterized tissue samples were collected from triple-negative breast cancer (TNBC) patients. MIBI-TOF images over 36 protein markers, but *M* = 24 markers were used in our training. A description of the technology and full list of these markers is given in S2.1.

#### 5.2.2 T-CyCIF

T-CyCIF (tissue cyclicimmunofluorescence) images of primary lung squamous cell carcinoma are represented in **T** ∈ℝ ^(*M,H*≈12000,*W* ≈14000)^. Segmentation patches are constructed in an similar fashion. The main salient difference from MIBI is the processed patch size **Y** ∈ℝ ^(*M*,128,128)^, and the segmentation patch size **Y** ∈ ℝ ^(*C*+1,128,128)^. This was intentionally done to demonstrate the scalability of CCIGAN. 44 markers were imaged in t-CyCIF, however we excluded background cell markers to yield *M* = 37 markers. A description of the technology and full list of these markers is given in the S2.2.

Unlike MIBI, a significant amount of data processing was done in order to analyze the data. Full treatment of data is given in S2.2.

#### 5.2.3 CODEX

CODEX images of colorectal cancer are represented in **T** ∈ℝ ^(*M*=10,1440,1920)^. These images are then further processed at a cell by cell basis into **Y** ∈ ℝ ^(*M*,64,64)^ patches. 10 markers relevant to tumor, immune, and muscle/epithelial cells were retained for training. More information on marker and segmentation preprocessing are found in S2.4.

### 5.3 Evaluation

To conduct fair experiments, all models were optimized, tuned, and set with similar parameters. They were also taken from their official online implementations and trained for 120 epochs or until convergence (max 150). CCIGAN is identical to our designed SPADE comparison baseline with the exception of the attention module.

#### 5.3.1 Image Evaluation and Reconstruction

First, we use the following evaluation metrics in order to compare with baseline results: adjusted *L*_1_ and MSE score, *L*_1_ and MSE score, structural similarity (SSIM) index (35) and cell based mutual information (MI) shown in Table 1. Bolded scores indicate the best scores. Equations and motivation are given in S3.

Three evaluation metrics were then used to conduct experiments and validate the trained model’s utility in generating biologically meaningful cellular proteins in the tumor microenvironment and ability to recapitulate and *quantify* previously established biological phenomena. Each subsection provides additional relevant information.

#### 5.3.2 Center of Mass (COM)

For a generated cell image, its weighted centroid, or center of mass, is the mean position of all the points in the cell weighted by a particular channel expression. Given a cell image ***X*** *∈*ℝ ^(*H,W*)^, with indices of the segmented cell *V ⊆*{1, …, *H*} × {1, …, *W*}, the COM 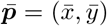 is defined as 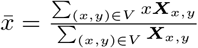 and 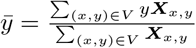.

For example, in the PD-1/PD-L1 experiment, we compute the COM of the CD8 T cell (cell of interest) weighted by PD-1 expression, given as 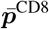, and the COM of all tumor cells weighted by PD-L1 expression, given as 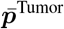. Since T cells located within the tumor microenvironment often have upregulated expression of PD-1, we assume that 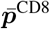 should have the roughly the same PD-L1 COM of all its surrounding tumor cells 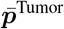. The center of mass score is defined below as the relative distance between 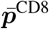 and 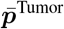, where *N* is defined as the number of patches:

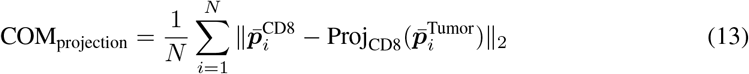

The projection function Proj(·) is used to project 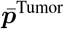 onto the CD8 T cell to ensure the expected COM of the tumor cells is inside of the CD8 T cell. As a reference we choose a random position 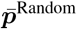 in the CD8 T cell (PD-1) which replaces 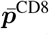 in Eq. 13 and compute the random COM score to show the effectiveness of the result. An example illustration is given in the figure in S3.1.

#### 5.3.3 Earth Mover’s Score

We develop Earth Mover’s (EM) Score to measure the dissimilarity between two distributions and the direction of distributional shift. While Earth Mover’s Distance is a nonnegative measurement of shift between two distributions, we incorporate the direction of shift in protein localization (positive indicating cell’s protein shifting towards another cell, negative otherwise) into our EM Score. For our Pan Keratin/CD8 experiment, we used EM to evaluate the shift in protein localization between CD8 T cells and tumor cells. EM can be generalized to experiments involving other cell-cell interactions, but for the purposes of clarity, below we describe our EM approach in the context of the Pan Keratin/CD8 experiment.

For a segmentation map, we add *T* tumor cells around one CD8 T cell. The COM for the *t*-th tumor (*t ∈{*1, …, *T*_*i*_ *}*) is defined as as 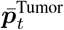. Similarly, the PD-1 COMs of the CD8 T cell by adding the *t*-th tumor is defined by 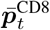. Initially when there are no tumor cells, 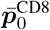 is the centroid of the CD8 T cell.

We proceed to define vector a vector ***v***_*t*_ which points from 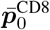, to the COM of the *t*-th tumor cell 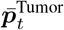. We also define vector ***u***_*t*_ which points from the previous COM 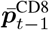 to the current COM 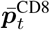 of the CD8 T cell. We define *θ*_*t*_ as the angle between ***v***_*t*_, ***u***_*t*_.

If cos *θ*_*t*_ *>* 0, that is to say if the cosine similarity is positive, the COM of a CD8 T cell 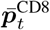, moves correctly towards the COM of the added tumor cell 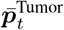. An illustration of the points and vectors is given in the figure in S3.2.

Formally:

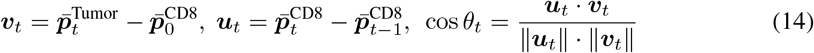

After obtaining the directional information, we use EM Score (36) to evaluate the changes in PD-1 expression of the CD8 T cell. Earth Mover’s Score, which measures the dissimilarity of two distributions, is used in this context to measure the protein localization shifts in PD-1 before and after adding a tumor cell. We consider each cell ***X*** in polar coordinates (*r, θ*) with respect to its centroid, integrate its expression along the radius coordinates, and evaluate the resulting histogram hist(***X***) along the angle coordinate. The 2nd figure in S3.2 shows an example histogram of cells by coordinate location.

This allows for the definition of distance for moving one histogram to another, i.e. 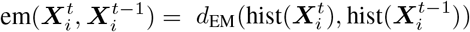, for the generated PD-1 expression of the CD8 T cell 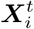 when adding the *t*-th tumor cell.

The final EM Score is defined as:

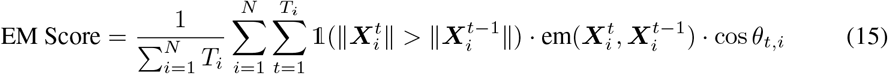

where the indicator function 1(·) = 1 if and only if 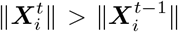, otherwise 1(·) = 0. This ensures that the biological constraint of PD-1 expression increasing as a response to added tumor cells is met. Recall, if cos *θ*_*t*_ *>* 0, 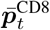 has moved in the direction of 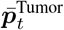, implying the shift in PD-1 expression is correct, and in turn increases the EM Score. By contrast, the EM Score decreases when 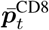 moves in the opposite direction.

This can be adapted and used for any two channel protein interactions (such as in the *β*-catenin and dsDNA experiment for analyzing Wnt pathways).

#### 5.3.4 Protein Expression and Cell Surface Area Trend Experiments

In this experiment, we used a Student’s *t*-test as the statistical hypothesis test to evaluate the correlations of the protein expression of a specified cell as a function of the area/number of surrounding cells in the specified cell’s microenvironment.

Given a generated protein channel ***X***_*i*_ ∈ℝ ^(*H,W*)^ and the segmentation map channel for the surrounding cells ***S***_*i*_ ∈ℝ ^(*H,W*)^, we compute the total area of the cells 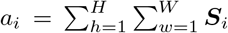, and the total expression level of the specified cell 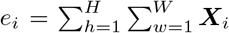. We then regress 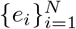 on 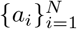 and assess significance of the slope using a *t*-test against the null of no change in expression as a function of surrounding cells.

Additional figures of other models and explanations for pan-keratin and CD8 are given in S3.4. Additional figures for t-CyCIF experiments are given in S3.6 and S3.7.

#### 5.3.5 Tumor Infiltrated and Compartmentalized Microenvironments

CCIGAN was used to compare protein localization in tumor infiltrated versus compartmentalized microenvironments. We used CCIGAN to predict on 200 (test data with 90-10 split) directly manipulated mixed and non-mixed tumor environment segmentation patches. For each experiment, we challenge the cell in question with an opposing cell in a microenvironment with macrophages (for example T Cell is challenged with Tumor cell to result in increased PD-1 expression in the T Cell) and use endothelial cells as control cells to show our result has biological significance. Similar to Online Methods 5.3.4‘s experimental settings, we compute the average expression of a specific marker for the cells of interest for all patches.

Data resulting from the experiment is located in S3.5. The increase in PD-L1 for the above tumor and macrophage scenarios (S3.5 table 7) indicate that CCIGAN has appropriately captured previously reported biological outcomes and is capable of quantifying these phenomena at single cell levels. Furthermore, the model is adaptable to various different types of tumor architecture depending on its training set to produce different hypothesis testing environments.

### 5.4 Search Algorithm

#### Search algorithm for detecting interactions

Here we provide a modular search algorithm framework to try to discover further cell-cell interactions in other channels. As a high level overview, the algorithm uses CCIGAN to automate and measure a specific cell’s change in a specified protein’s expression level due to user specified microenvironment changes. The algorithm allows a user to change and specify such changes as cell type, shape, size, and quantity. If the change in expression is greater than a user specified input, then the particular instance is logged. It is important to note that this tool is meant to guide and search for particular interesting interactions and still susceptible to issues such as noisy segmentations.

##### Algorithm 1 Search Algorithm

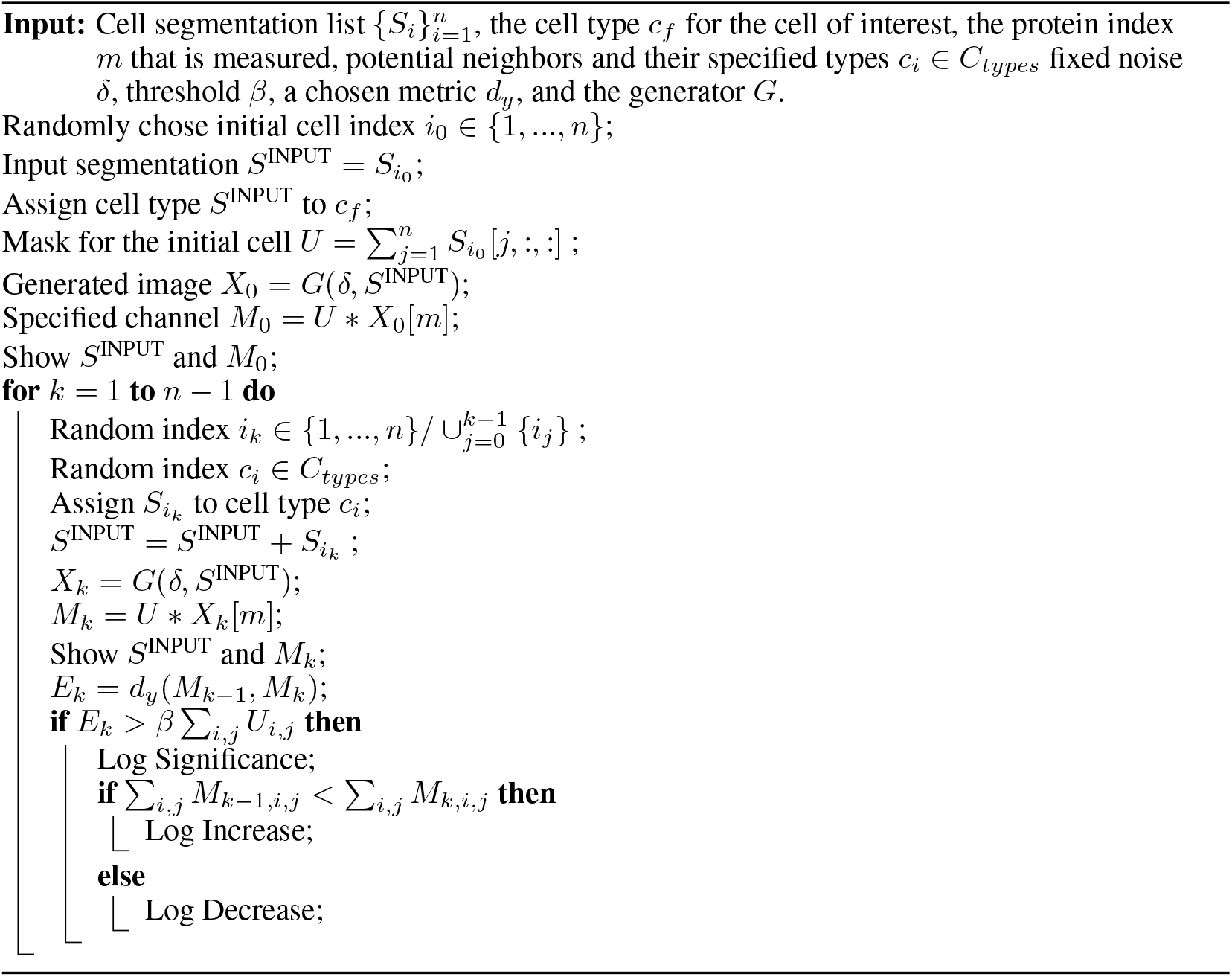

## Supplementary Materials

### 1 Model

#### 1.1 Model Architecture

The detailed architecture of our generator is shown on Table 3.

**Table 3:** Architecture details of CCIGAN’s generator

| Layers | Output Size | Generator |
| --- | --- | --- |
| Linear | (128, 4, 4) | Linear $128 \times 2048$ |
| Upsampling | (128, 8, 8) | Upsampling $2 \times 2$ |
| SPADE ResBlk-1 | (128, 8, 8) | SPADE 128, Leaky ReLU<br>Convolution $3 \times 3$<br>SPADE 128, Leaky ReLU<br>Convolution $3 \times 3$ |
| Upsampling | (128, 16, 16) | Upsampling $2 \times 2$ |
| SPADE ResBlk-2 | (128, 16, 16) | SPADE 128, Leaky ReLU<br>Convolution $3 \times 3$<br>SPADE 128, Leaky ReLU<br>Convolution $3 \times 3$ |
| Upsampling | (128, 32, 32) | Upsampling $2 \times 2$ |
| SPADE ResBlk-3 | (64, 32, 32) | SPADE 128, Leaky ReLU<br>Convolution $3 \times 3$<br>SPADE 64, Leaky ReLU<br>Convolution $3 \times 3$<br>SPADE 64, Leaky ReLU<br>Shortcut Convolution $3 \times 3$ |
| Upsampling | (64, 64, 64) | Upsampling $2 \times 2$ |
| SPADE ResBlk-4 | (64, 64, 64) | SPADE 64, Leaky ReLU<br>Convolution $3 \times 3$<br>SPADE 64, Leaky ReLU<br>Convolution $3 \times 3$ |
| Convolution | (24, 64, 64) | Leaky ReLU, Convolution $5 \times 5$ |
| Convolution | (24 * 8, 64, 64) | Leaky ReLU, Convolution $5 \times 5$ |
| Group SPADE | (24 * 8, 64, 64) | [SPADE 8] * 24 |
| Modulation | (24 * 64, 64, 64)<br>(24, 64, 64) | [Outer Product $8 \otimes 8$] * 24<br>Convolution $1 \times 1$, Sigmoid |
| Output | (24, 64, 64) | Sum residual, Sigmoid |

where ResBlk is the residual block with skip connection used in ResNet (37), and SPADE is the spatially-adaptive normalization layer. The detailed architecture of our discriminator is shown on Table 4.

**Table 4:**
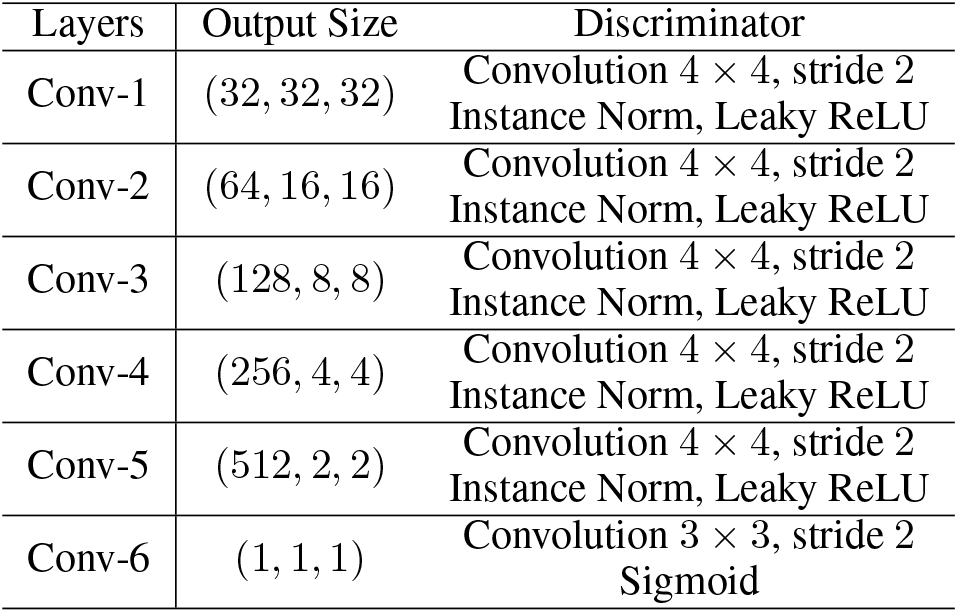
Architecture details of CCIGAN’s discriminator

#### 1.2 Implementation Details and Training Regimen

Our implementation of the generator applies Spectral Norm to all layers (38). The discriminator’s input is the output of the generator concatenated with the segmentation patch [**X, S**] and [**Y, S**] for the ground truth. Finally CCIGAN uses ADAM (*lr*_*G*_ = 0.0004, *lr*_*D*_ = 0.0001) with GAN loss and feature matching loss. Full training details and loss functions are as follows.

#### 1.3 Model Training

*G* is the generator and *D* is the discriminator for CCIGAN. Given segmentation map **S**, ground truth **Y** and noise *δ*, the generated image is **X** = *G*(**S**, *δ*). The input of the discriminator is the cell image conditioned on the segmentation map **S**. We use LSGAN loss (39) in CCIGAN, which is defined as follows:

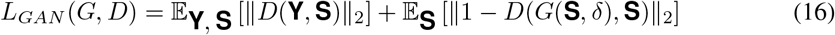

In addition to GAN loss, we also use feature matching loss (24) during training expressed as:

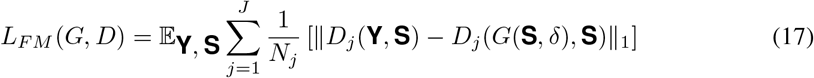

where *D*_*j*_ is *j*-th layer feature map of the discriminator for *j ∈*{1, …, *J*}, and *N*_*j*_ is the number of elements in *j*-th layer. Consequently, the objective function for training is given as follows:

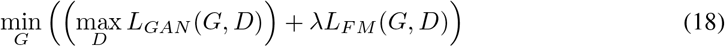

where *λ* = 10. Due to the size of cell patch is (64, 64), we do not use multi-scale discriminators and perceptual loss in CCIGAN and other baseline models e.g. SPADE and pix2pixHD.

In training, we use ADAM as the optimizer. The generator learning rate is *lr*_*G*_ = 0.0004 and the discriminator learning rate is *lr*_*D*_ = 0.0001. We train CCIGAN 120 epochs with a training set of 5648 cell patches. We train other baseline models for 120 epochs or until they converge (max 150). The full details of training of CCIGAN and baselines are shown as Table 5. The hyperparameters of each model are fine-tuned to get better performance. The training time was roughly equal for all models. In particular, CCIGAN was around 1.2 times slower than the SPADE baseline on a single Tesla V100 GPU.

**Table 5:** Hyperparameters of models

| Metrics | Ours | SPADE | Pix2PixHD | CycleGAN |
| --- | --- | --- | --- | --- |
| $lr_G$ | 0.0004 | 0.0008 | 0.0002 | 0.0002 |
| $lr_D$ | 0.0001 | 0.0001 | 0.0002 | 0.0002 |

#### 1.4 Model Interpretability and Generativeness

Examining the model’s persistent vectors ***s***_*m*_, we can try to understand if there is a match between real world protein markers and the representations of ***s***_*m*_. For example, the vector ***s***_pk_ for pankeratin attends to tumor cells and ***s***_CD8_ attends to CD8 T cells at pixel pairwise levels. It follows that in a simple experiment where corresponding ***s***_CD8_ ↔***s***_pk_ vectors are exchanged internally in the attention module (Eq. 10, Fig. 1D (Step 3), outer product) we may observe a lower expression for tumor cells in channel *m*_pk_ and a lower expression for CD8 T cells in channel *m*_CD8_ since tumor cells do not express CD8 and CD8 T cells do not express pan-keratin. As a control, we also switch surface membrane markers HLA Class 1 and dsDNA markers as they are present in all cells and have very similar average expression values (***s***_HLAc1_ ↔ ***s***_dsDNA_). Accordingly, for our control, we expect to see negligble changes. We define the expression ratio as 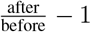.

**Table 6:** *s*_*m*_ persistent vector interpretability experiments.

| Protein Markers | CD8 | pan keratin | HLA Class 1 | dsDNA |
| --- | --- | --- | --- | --- |
| Expression Ratios | -0.373 | -0.145 | -0.054 | -0.0012 |

In Table 12, we can see a larger magnitude decrease of the expression ratios in the ***s***_CD8_ ↔***s***_pk_ experiment and a minute difference in the ***s***_HLAc1_ ↔***s***_dsDNA_. Further visualizations (Fig. 6) and discussion (model generativeness, Fig. 7) are as follows.

**Figure 6:**
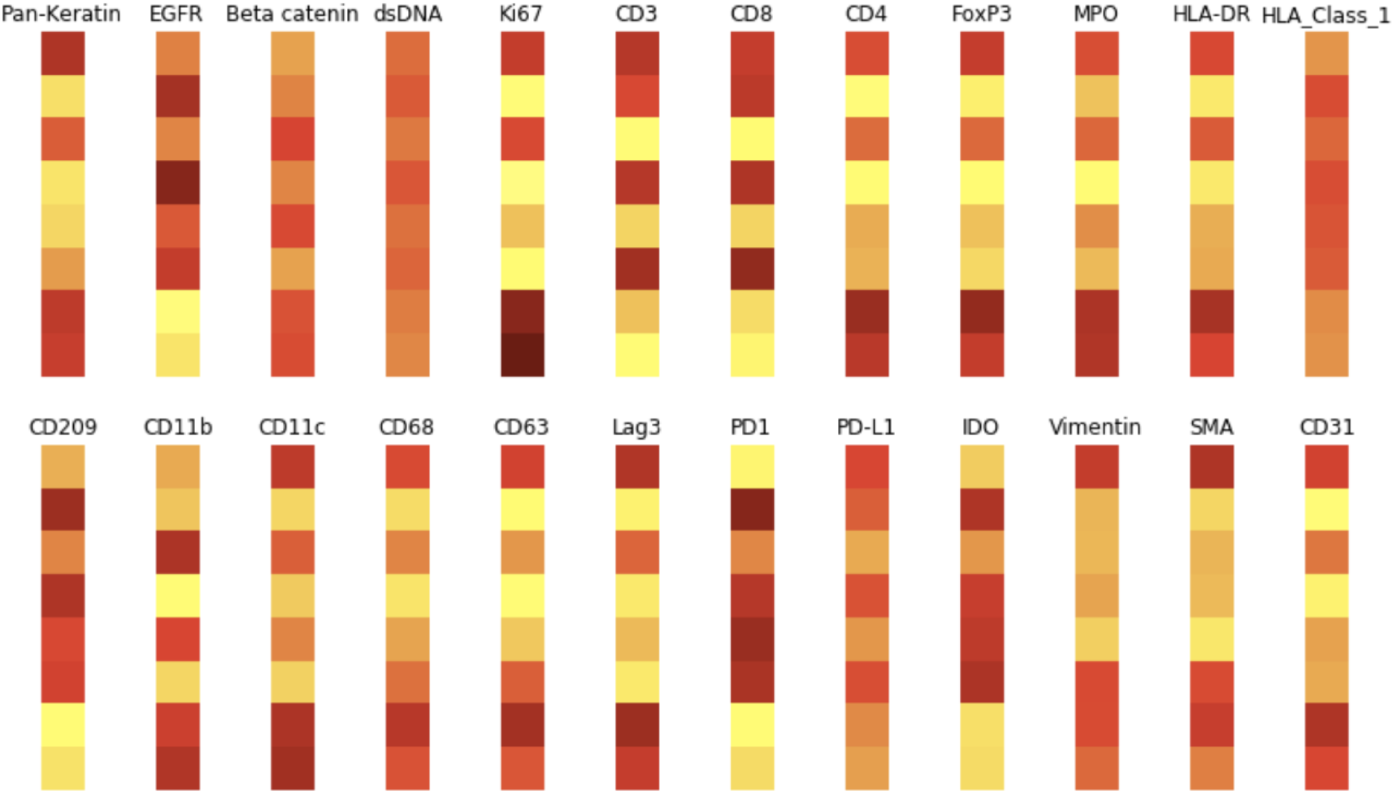
Persistent vectors *s* for various channels.

**Figure 7:**
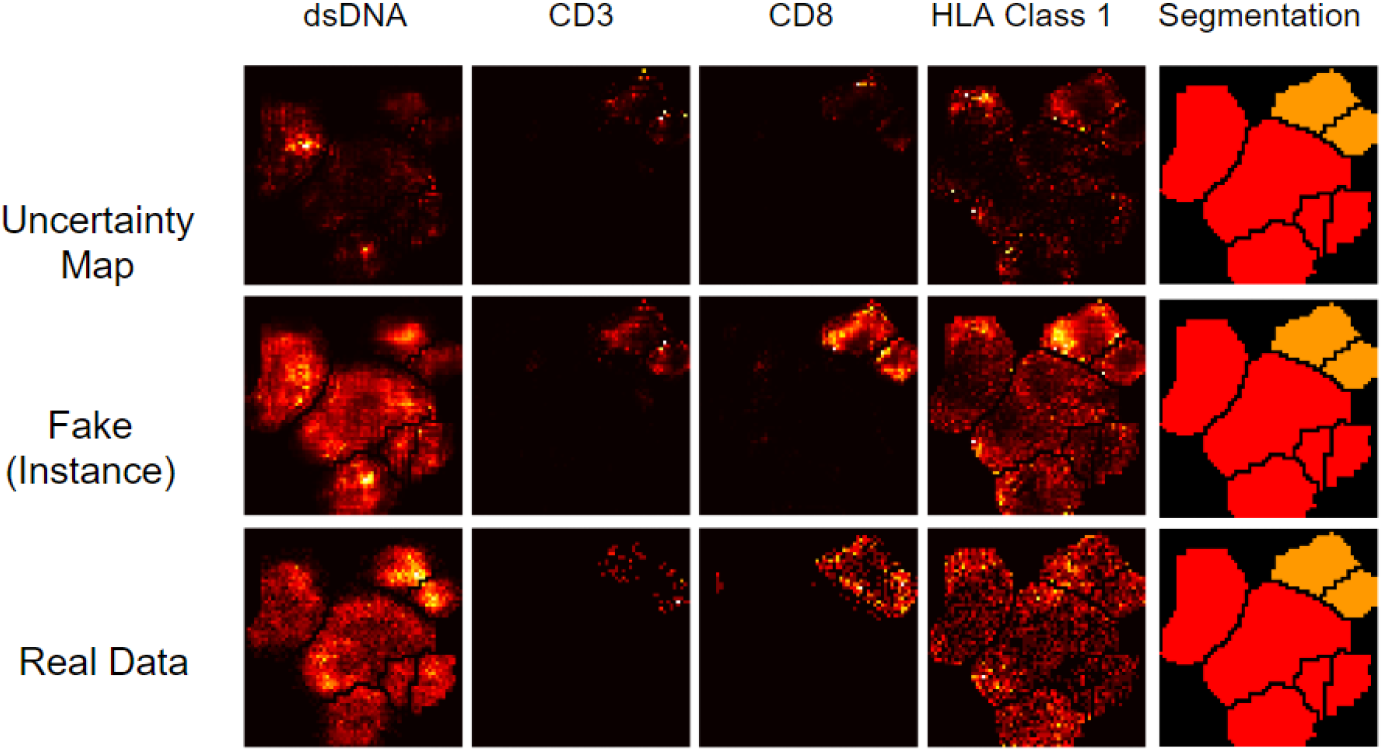
Uncertainty maps illustrating model generativeness.

**Figure 8:**
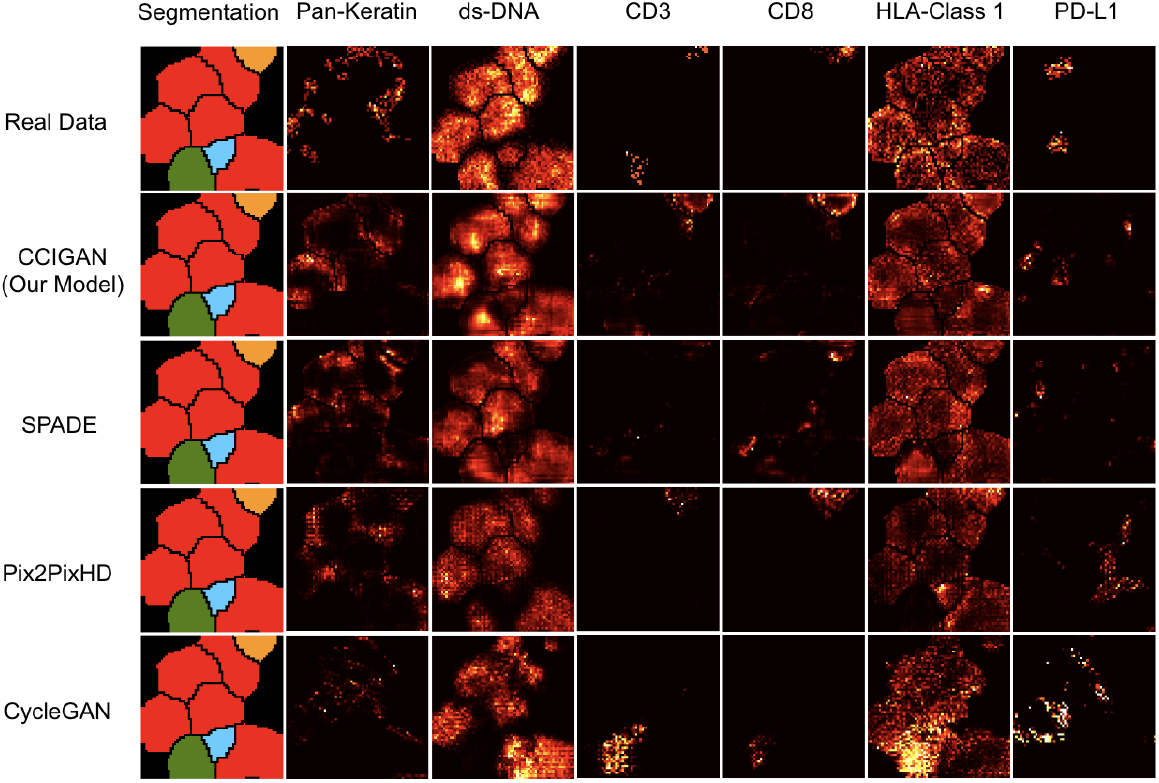
Examples generated from a segmentation for certain channels for different models. The segmentation patch is the one hot encoded patch collapsed and colored into 1 channel. The horizontal labels represent protein markers and the vertical labels are each of the generative models.

Fig. 6 shows the persistent vectors ***s***_*i*_ for all proteins. Note the similarity between CD3 and CD8 T cell protein markers and the similarity between dsDNA and HLA Class 1 surface membrane proteins (expressed in all cells). It is also important to make the distinction that sparse markers (while different) are similar in state. This is due to the lack of training data for rare cell types, making it difficult for the model to reason on such a small sample size.

Fig. 7 shows the generativeness of CCIGAN through an uncertainty map over 100 instances (random noise). An uncertainty map shows the differences per pixel (*x, y*) location. The higher intensities indicate a higher probability of changing at the specified (*x, y*) location.

### 2 Data and Data Processing

#### 2.1 MIBI-TOF

Multiplexed ion beam imaging by time-of-flight (MIBI-TOF) represents a novel technology that can accurately quantify and spatially resolve cellular protein expressions at the single cell level within tissue samples. Given a tissue sample that is first stained with protein-specific antibodies tethered to elemental metals, MIBI-TOF bombards the sample with atomic ions (i.e. 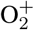) from a primary ion beam. This causes the release of elemental isotopes and tissue-specific material which can be quantified in a mass spectrometer (40).

For MIBI, the markers used (total 24) in our experiments are: Pan-Keratin, EGFR, Beta catenin, dsDNA, Ki67, CD3, CD8, CD4, FoxP3, MPO, HLA-DR, HLA-Class-1, CD209, CD11b, CD11c, CD68, CD63, Lag3, PD1, PD-L1, IDO, Vimentin, SMA, CD31. The markers we didn’t use (total 12) in our experiments are: CD16, B7H3, CD45, CD45RO, Keratin17, CD20, CD163, CD56, Keratin6, CSF-1R, p53, CD138. These were not used primarily due to blank expressions.

#### 2.2 T-CyCIF

The t-CyCIF images are obtained through iterative cycles of incubating the sample with antibodies (markers) conjugated directly with flurophores that will bind to a specific protein of interest, imaging the samples with a microscope to record the light emitted at each location of the protein, then deactivating the fluorescence signal by soaking the sample in a compound (4). At each iteration, different antibodies that bind to different proteins of interests are applied. After obtaining multiple images with few channels at each iteration, the images are stitched together to produce a high-dimension image.

For t-CyCIF, the markers used (total 37) are: DAPI1, DAPI2, DAPI3, LAG3, ARL13B, DAPI4, KI67, KERATIN, PD1, DAPI5, CD45RB, CD3D, PDL1, DAPI6, CD4, CD45, CD8A, DAPI7, CD163, CD68, CD14, DAPI8, CD11B, FOXP3, CD21, DAPI9, IBA1, ASMA, CD20, DAPI10, CD19, GFAP, GTUBULIN, DAPI11, LAMINAC, BANF1, LAMINB. The markers were didn’t use (total 7) are: A488background1, A555background1, A647background1, A488background2, A555background2, A647background2, A488background3.

##### 2.2.1 T-CyCIF Data Preprocessing

For t-CyCIF data, each cell was first clustered to one of many cell types. Using cell features (log cell expression data), we clustered the cells using the 26 non-DAPI markers. We exponentiated the cell features data to restore the data to raw cell intensities, then quantile clipped each marker to retain only the data from 1% to 99.5%. After 0-1 min-max rescaling the data, we clustered the cells using the Phenograph tool, developed in the Dana Pe’er Lab.

Let *N* denote the nuclear probability matrix of the image where *N*_*i,j*_ = Pr[nucleus at index (*i, j*)] and let *N ′* be the 0-1 nuclear segmentation mask, 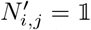 [cell nucleus at index (*i, j*)]. Cytoplasm probabilities were also provided, the cytoplasm probability mask is denoted as *C*, where *C*_*i,j*_ = Pr[cytoplasm at index (*i, j*)]. *N, N ′, C* were provided to us from (27). We modified the cytoplasm probability mask to a 0-1 cytoplasm mask *C′* where 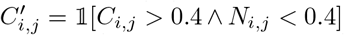 by choosing these thresholds ourselves. Then we overlayed the nuclear and cytoplasm masks together to form a mask *T* of the total cell, where *T*_*i,j*_ = 1 [a cell occurs at index (*i, j*)]. Because there are small holes in this mask, we filled the holes using SK-image’s morphology tools. To yield the final segmentation mask *F*, we dilated the nuclear segmentation mask *N ′* with 2 iterations and perform an element-wise mask with mask *T*, where 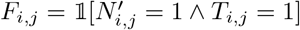.

##### 2.2.2 T-CyCIF Data Split

We split the t-CyCIF primary lung cancer dataset into four segments to test our hypothesis that CCIGAN can learn biological relationships specific to tumor microenvironments with varying characteristics. Three segments (highly, semi, and poorly infiltrated) were each manually selected based on the visible proportion of lymphocyte presence within the tumor microenvironment. Figures 9 and 10 show the segments in relation to the lymphocyte and keratin expression. More detailed descriptions of each individual dataset segment are as follows.

**Figure 9:**
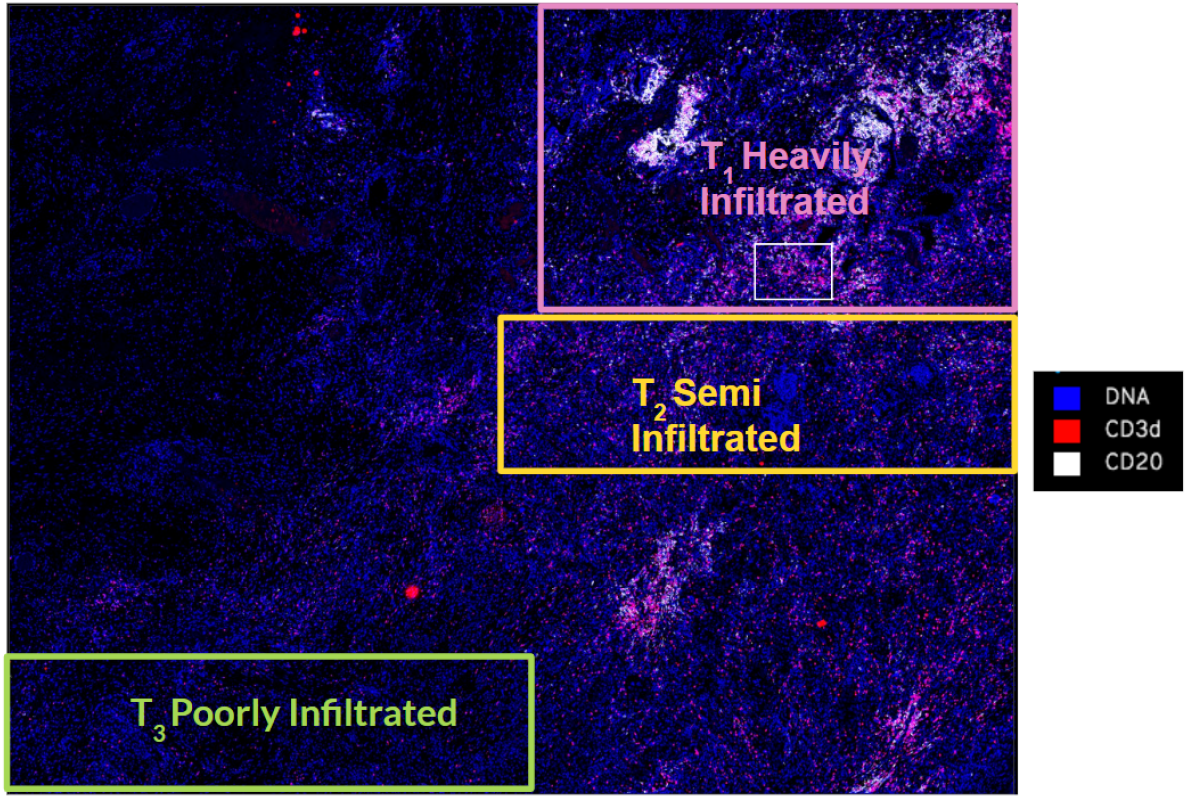
Splits of the tissue with lymphocyte cell protein markers highlighted

**Figure 10:**
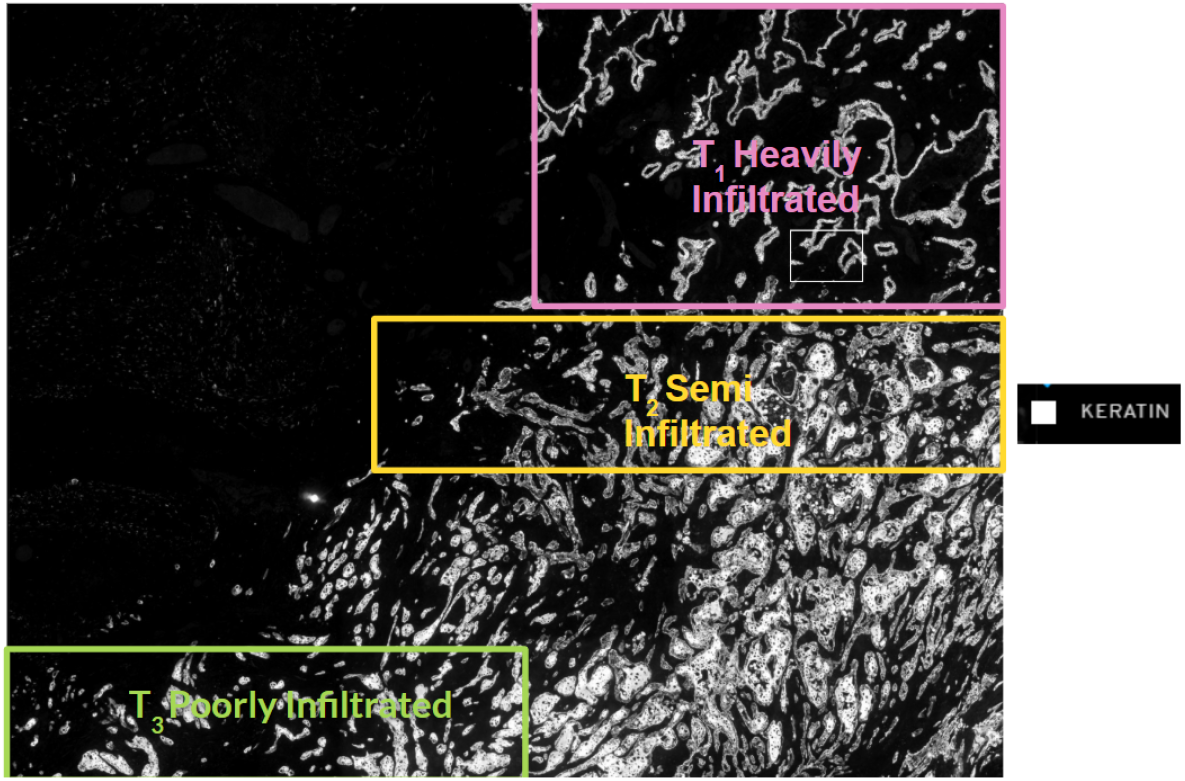
Splits of the tissue with keratin positive tumor cell type highlighted

1. *T*_1_ Infiltrated - High proportion of lymphocyte presence within the tumor microenvironment. Indicative of a strong inflammatory anti-tumor immune response.
2. *T*_2_ Semi-infiltrated - Medium levels of lymphocyte infiltration within the tumor microenvironment.
3. *T*_3_ Poorly-infiltrated - Few lymphocytes present within the tumor microenvironment indicative of a poor anti-tumor immune response.

#### 2.3 CODEX

We trained CCIGAN on 6 patients in the DII cohort of the colorectal cancer CODEX data (3) with high frequency of ICOS+CD8 T cells and ICOS+Tregs (image no. 21, 22, 28, 44, 54, 59). We used 10 markers total: CD44 - stroma, CDX2 - intestinal epithelia, CD8 - cytotoxic T cells, PD-L1 - checkpoint, Ki67 - proliferation, CD4 - T helper cells, aSMA - smooth muscle, PD-1 - checkpoint, ICOS - costimulator, CD31 - vasculature. We also grouped cell types in 11 groups: ‘tumor cells’, ‘CD68+ macrophages’, ‘smooth muscle’, ‘stroma’, ‘CD4+ T cells’, ‘CD8+ T cells’, ‘vasculature’, ‘immune cells’, ‘Tregs’, ‘undefined’.

Segmentations were obtained from the CODEX image processing software and segmentations from multiple Z axes reported in the cell typing CSV file from Schürch et al. (3) were joined together to form one segmentation per patient image. More details on data preprocessing are in the code provided.

#### 2.4 Data Limitations

Examples are given in Fig. 11.

**Figure 11:**
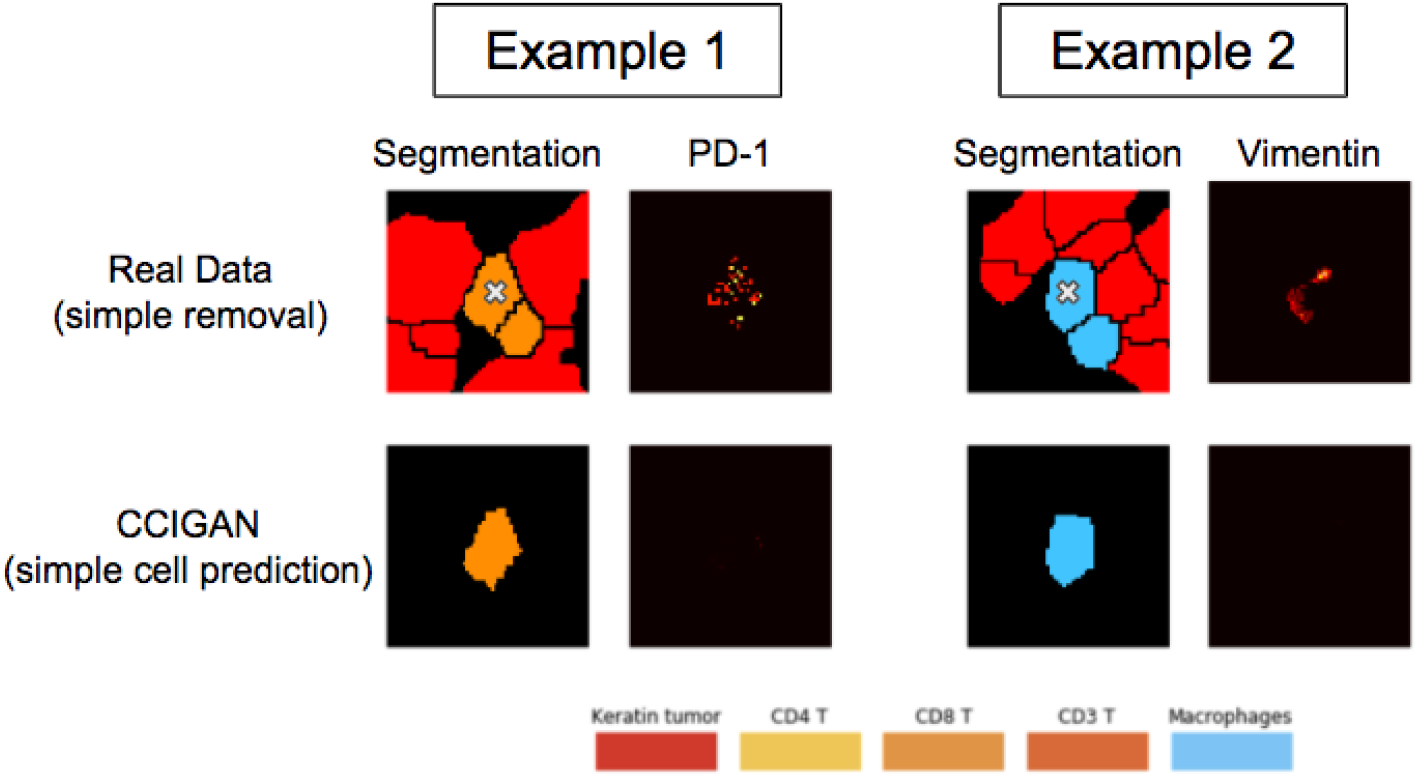
Data limitations regarding manually isolating cells and their protein localizations. Simply deleting cells surrounding a target cell does not reveal the true protein localization of the target cell (with the X marker), as the protein localization will still assume a neighboring interactions. In Example 1, the target CD8 T cell should only express PD-1 when it is surrounded by PD-L1 expressing tumor cells which upregulate the PD-1 expression. Note in the real data how deleting the surrounding cells do not change its PD-1 protein localization. CCIGAN learns that for an isolated CD8 T cell, PD-1 is not upregulated and therefore not expressed.

### 3 Evaluation

#### 3.1 Center of Mass (COM)

Fig. 12 shows an illustration of the center of mass and projected center of mass of a patch of tumor and CD8 T cells.

**Figure 12:**
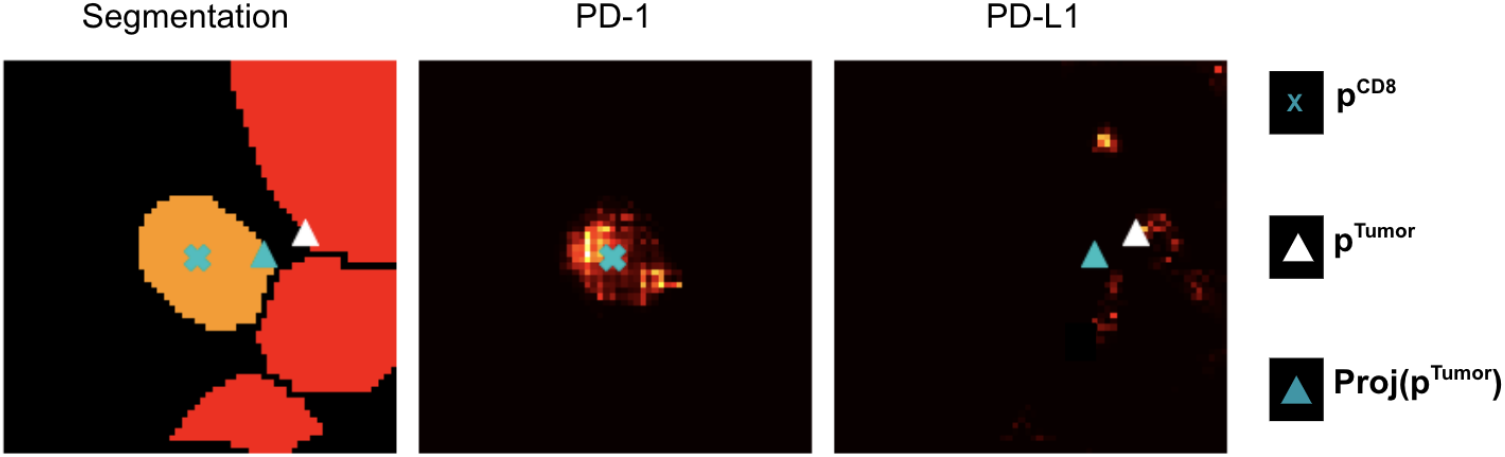
An example illustration of the center of mass (COM) nomenclature. Note the projection onto the CD8 T cell. This provides a more consistent measurement across different patches by projecting ***p***^Tumor^ onto the CD8 T cell.

#### 3.2 Earth Mover’s Score

Figures 13 and 14 demonstrate the directional and histogram shifts as a function of adding more tumor cells.

**Figure 13:**
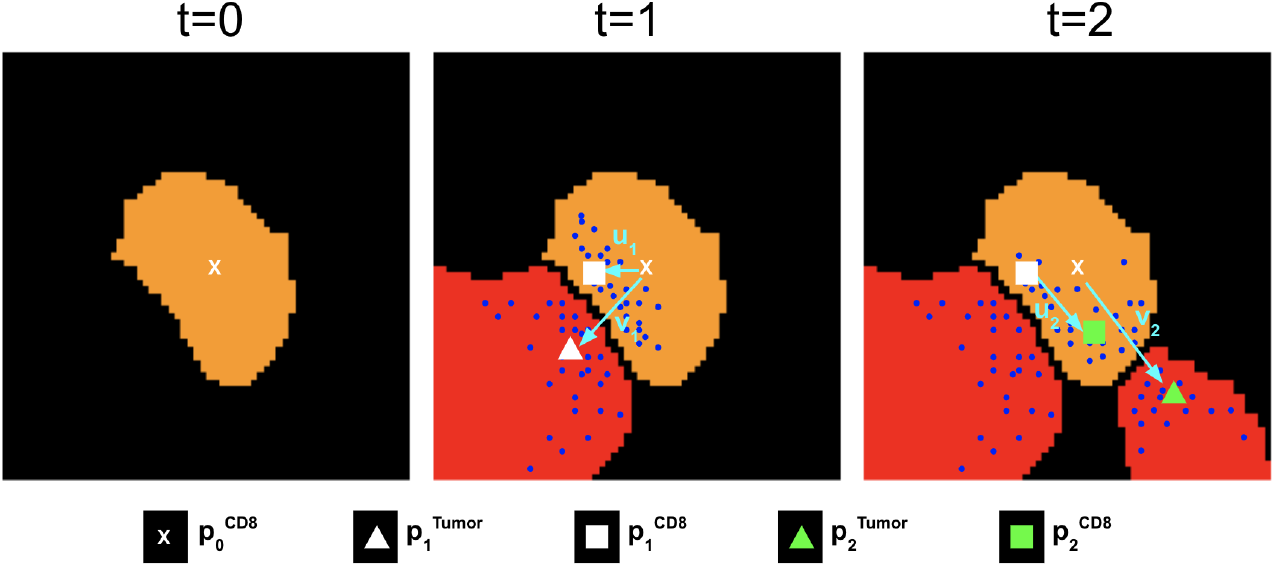
An example illustration of the points and vector nomenclature from Online Methods 5.3.3. The blue dots are the expression of PD-1 and PD-L1 proteins. The cyan arrows show the vectors ***v***_*t*_ and ***u***_*t*_. Note the shift in expression of the PD-1 as a response to the added tumor’s PD-L1 expression.

**Figure 14:**
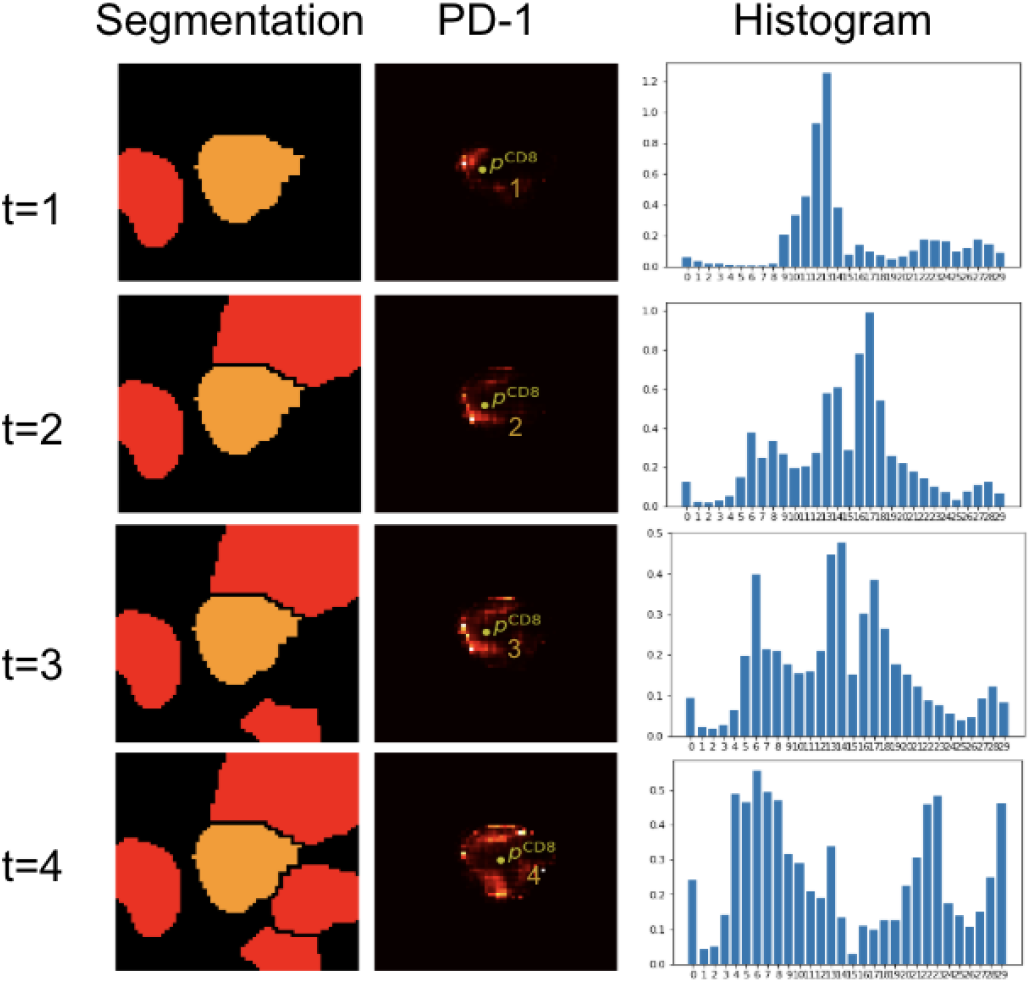
Example illustration of how a CD8 T cell’s (orange) PD-1 histogram changes as a function of iteratively added tumor cells.

#### 3.3 Reconstruction Metrics

Given the generated image set 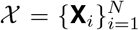 and the ground truth set 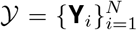 with **X**_*i*_, **Y**_*i*_ ∈

ℝ ^(*M,H,W*)^, the *L*_1_/MSE score is defined as follows,

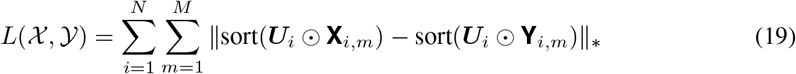

where ‖· ‖_∗_ can be either *L*_1_ or *L*_2_ norm, ⊙is the element-wise product, **X**_*i,m*_ and **Y**_*i,m*_ are the *m*-th channel of the *i*-th cell patch, ***U***_*i*_ ∈ {0, 1}^(*H,W*)^ is the mask matrix which masks all the cells in *i*-th patch. For any matrix ***A***, sort(***A***) is the sort function that sorts all entries of ***A***. The sorting function ensures our metrics are position independent and only measures the intensity of the generated image and ground truth. The score function *L*(𝒳, *𝒴*) only computes the loss of sorted expression inside of the cells. Then we add penalization for expression outside of cells. The adjusted *L*_1_/MSE score is introduced as follows,

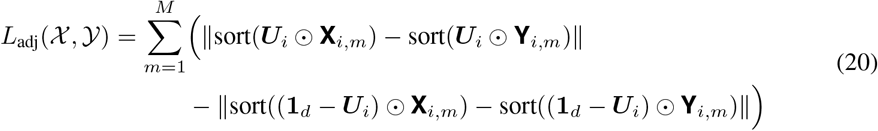

where **1**_*d*_ is the matrix with all entries equal to 1. A smaller score indicates a better result. For any two images ***X, Y*** ∈ [0, 1]^(*H,W*)^, the SSIM and MI are defined as:

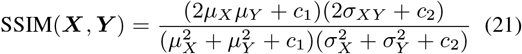

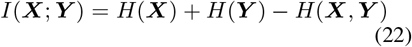

where *H*(·) is entropy, *µ*_*X*_ and *σ*_*X*_ are the mean and standard deviation of **X**, *c*_1_, *c*_2_ are constants. Then the SSIM between 𝒳, 𝒴 is

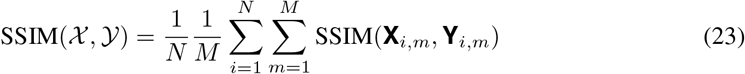

In cell based MI, test patches are processed at a cell-cell basis where their mutual information is computed with the corresponding cell in the ground truth. For the generated image **X**_*i*_ of the *i*-th patch, we assume there are *T*_*i*_ cells in the *i*-th patch. Then for each cell *t*, the pixels of *m*-th channel of the *t*-th cell in the *i*-th patch can be expressed as a vector 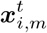. Hence, the cell based MI is formulated as:

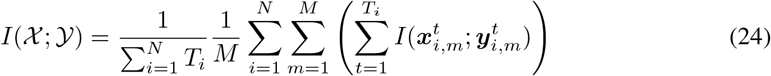

The SSIM measures the similarity between the generated image and the ground truth. For SSIM, we use HLA Class 1 and dsDNA due to the their expressions in all cells. If all channels were considered, the SSIM would be uninformative due to the majority of the channels being blank or sparse. The MI measures the information shared between generated image and ground truth at a cell by cell basis where we consider all channels. Consider the example where a model generates no expression in marker *m* but the real data has expression in *m*, the MI would be 0 and vice versa. Higher SSIM and MI values indicate better results.

#### 3.4 Pan-keratin and CD8 Expression

The pan-keratin/CD8 experiment is similar to Fig. 23‘s orientation except the center cell (cell of interest) is a tumor cell (red) and the adjacent neighboring cells are CD8 T cells (orange). CCIGAN predicted a decrease in tumor cell pan-keratin expression with respect to increasing CD8 T cell area/number (Fig. 15). This is juxtaposed to the tumor cell control where there is no change in the pan-keratin level as the number of neighboring tumor cells is increased.

**Figure 15:**
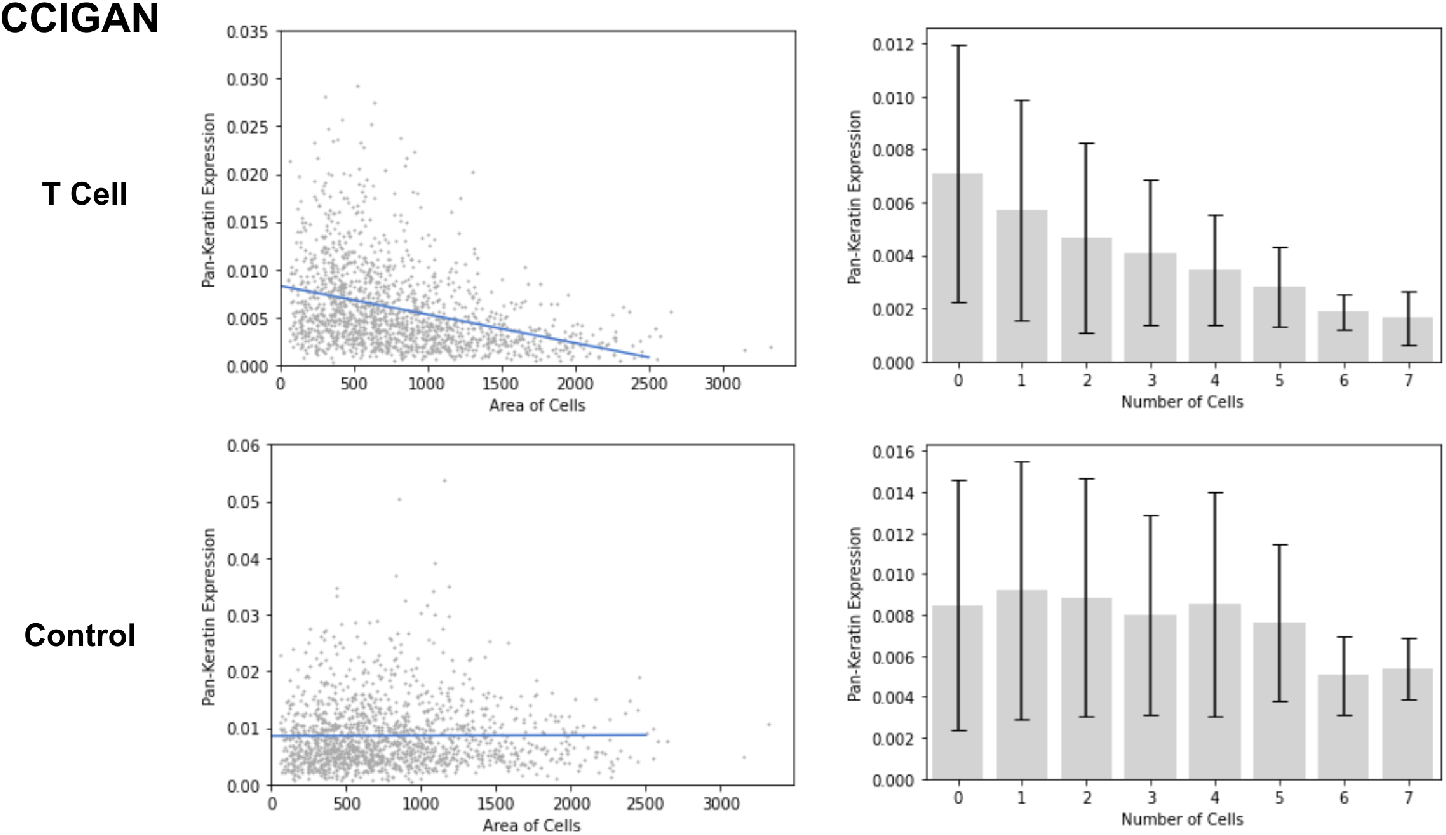
CCIGAN experiment for adding CD8 T cells and tumor cells (control) around a tumor cell.

SPADE does not predict a decrease in tumor cell pan-keratin expression with respect to increasing CD8 T cell area/number and shows no difference in pan-keratin expression trends between the T cell and control groups (Fig. 16).

**Figure 16:**
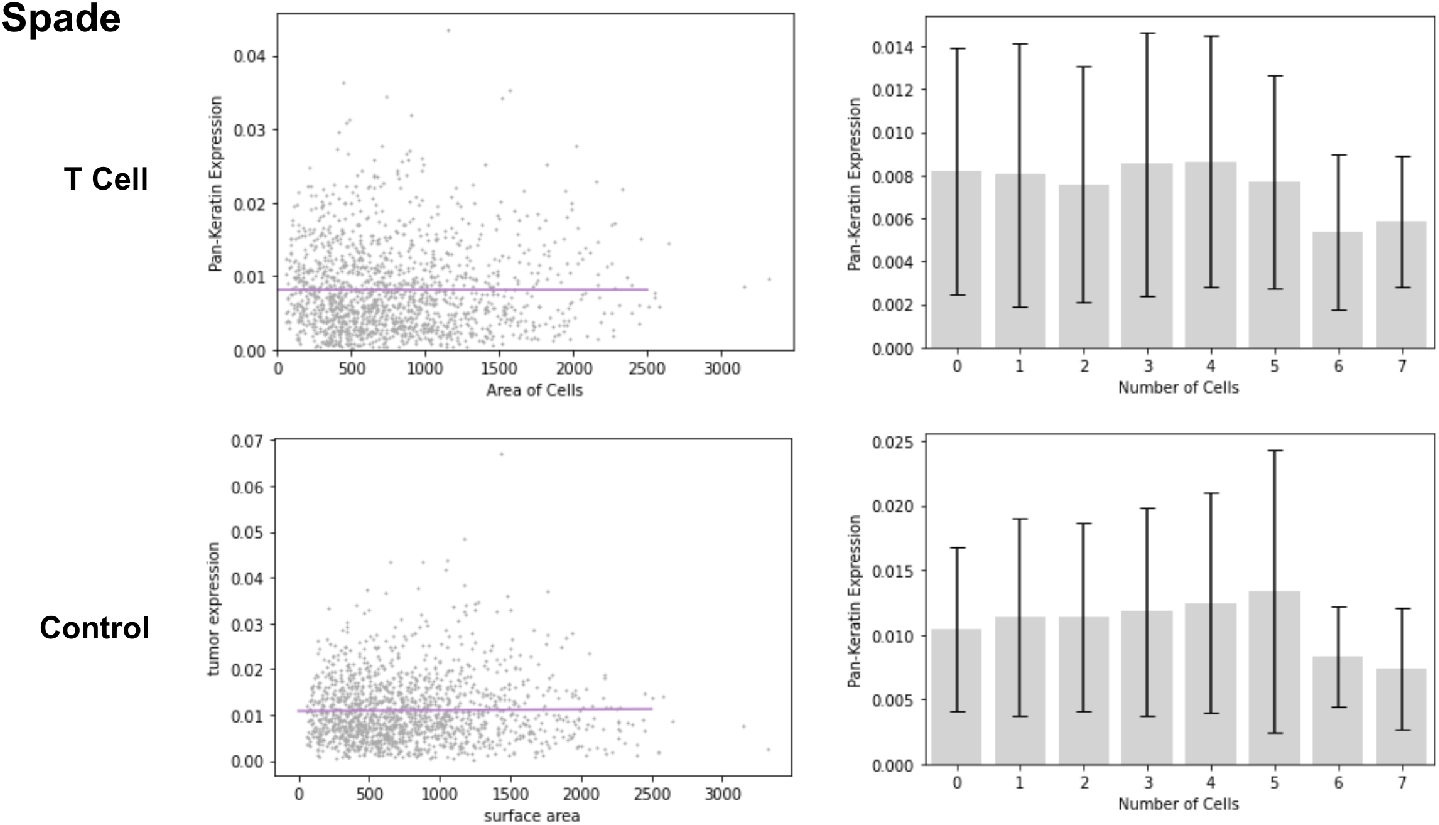
SPADE experiment for adding CD8 T cells and tumor cells (control) around a tumor cell.

pix2pixHD erroneously predicts an increase in tumor cell pan-keratin expression with respect to increasing CD8 T cell area/number and shows no difference in pan-keratin expression trends between the T cell and control groups (Fig. 17).

**Figure 17:**
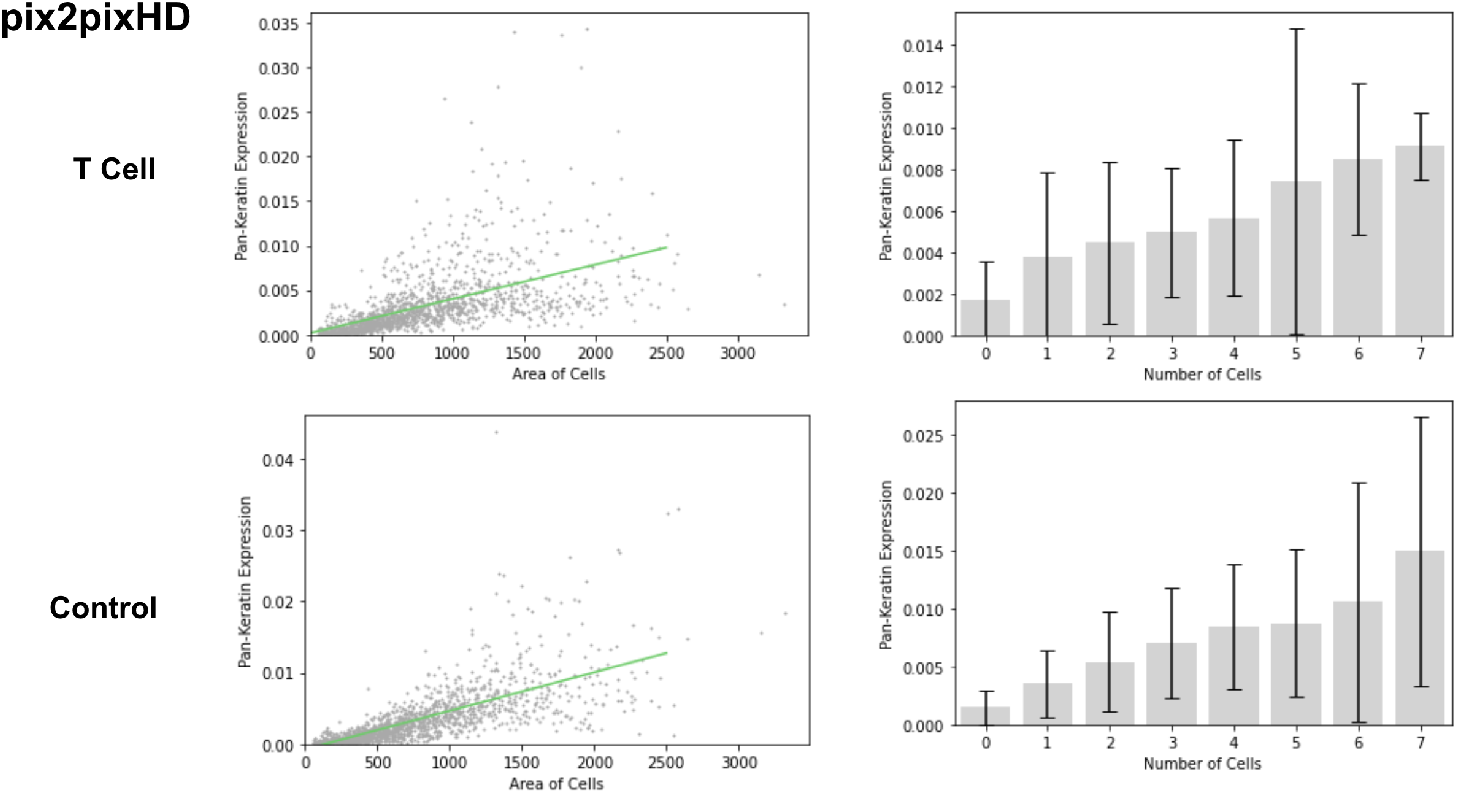
pix2pixHD experiment for adding CD8 T cells and tumor cells (control) around a tumor cell.

CycleGAN fails to predict a decrease in tumor cell pan-keratin expression with respect to increasing CD8 T cell area/number and shows no difference in pan-keratin expression trends between the T cell and control groups (Fig. 18).

**Figure 18:**
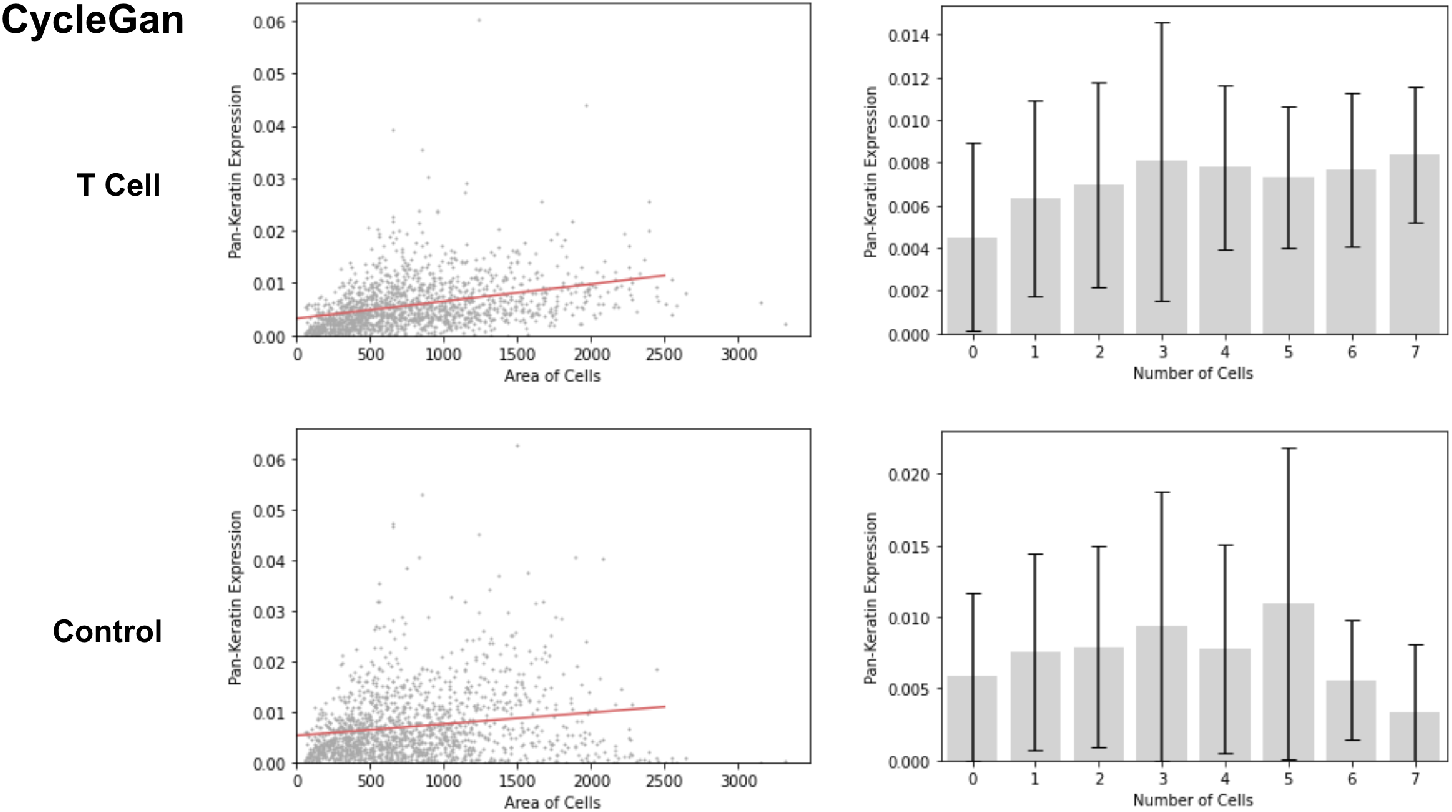
CycleGAN experiment for adding CD8 T cells and tumor cells (control) around a tumor cell.

**Figure 19:**
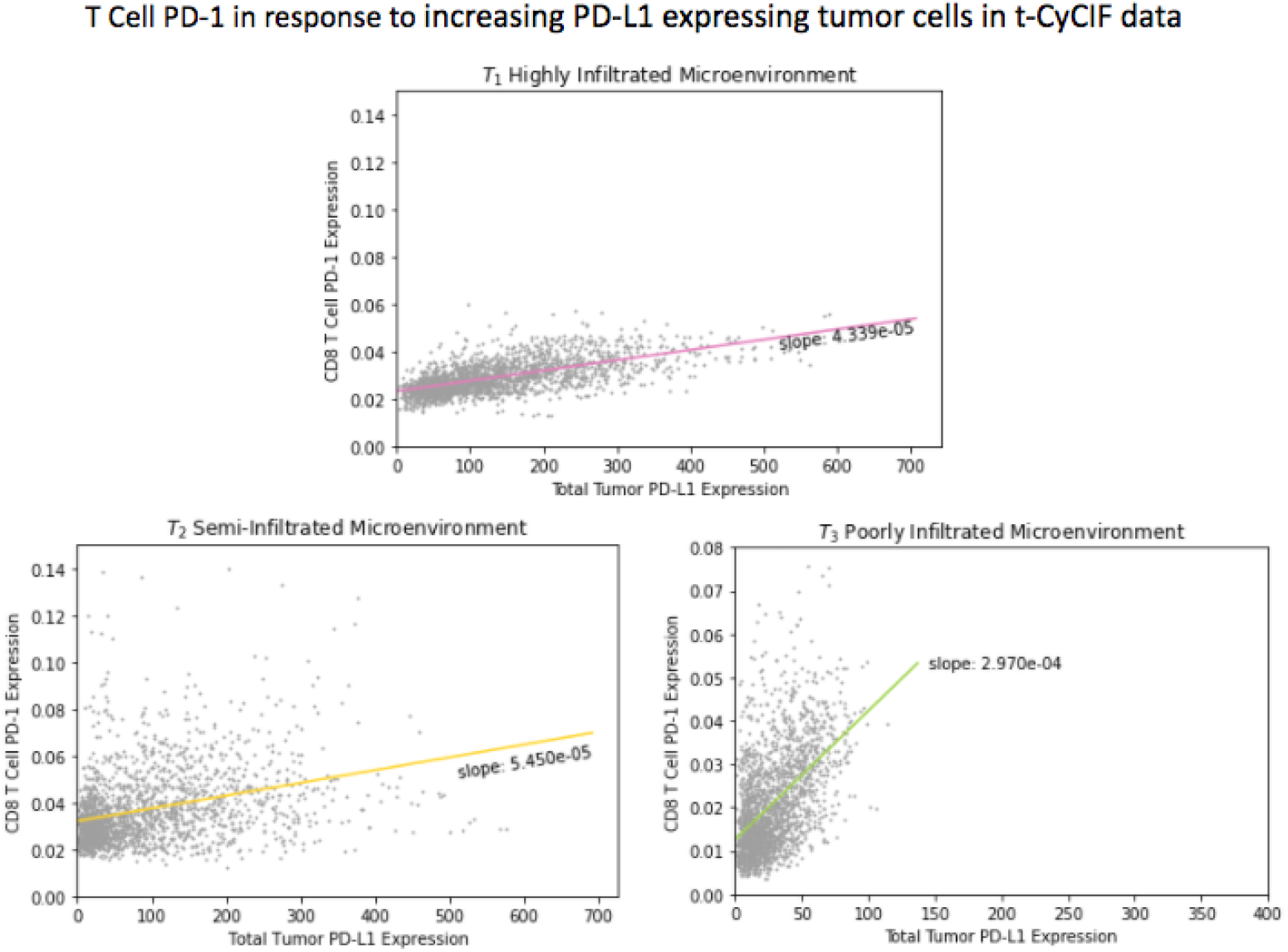
PD-1/PD-L1 experiment on tumor microenvironments with varying levels of tumorinfiltration in t-CyCIF (S3.6) with respect to total surrounding PD-L1 expression.

#### 3.5 Tumor Infiltrated and Compartmentalized Microenvironments

Keren et al. (2018) determined that in situations of mixed tumor-immune environments, where immune cells freely infiltrated the tumor, the tumor cells predominantly expressed PD-L1. Conversely, in situations of compartmentalized tumors, where there is a greater degree of physical separation between immune and tumor cells, macrophages were the predominant source of expressed PD-L1, particularly at the tumor boundary.

These findings were recapitulated by CCIGAN. For a patient with a mixed tumor environment, when trained with mixed patient samples, CCIGAN reported increased PD-L1 expression on tumor cells. Furthermore, CCIGAN was able to quantify this difference in expression at the single cell level, reporting a tumor to macrophage PD-L1 expression ratio of approximately 3.2 and 1.75 for patients A and B respectively.

Conversely, when trained with compartmentalized patient samples, CCIGAN reported increased PD-L1 expression on macrophages adjacent to tumor cells when compared to macrophages adjacent to normal endothelial (inert) cells. This difference was quantified as a ratio of PD-L1 expression of tumor-adjacent macrophages to endothelial-adjacent macrophages, approximately 1.85 and 2.7 for patient C and patient D respectively.

Below are tables of the data used to generate Fig. 4B of the main paper. Results from testing increased PD-1/PD-L1 expression from the bolded cell being challenged with another cell type in its microenvironment are located in table 7. The 3rd column shows summed pixel intensity of the specified protein expression.

In table 8, the second row shows that even when using the trained compartmentalized model to predict on mixed segmentation patches, CCIGAN still reports a 26% (patient C, 0.00408/0.00324) and 19% (patient D, 0.00608/0.00510) increase of macrophage PD-L1 expression in the compartmentalized compared to mixed microenvironment (Table 8). This observation of higher macrophage PD-L1 expression in the compartmentalized environment supports Keren et al.‘s findings.

**Table 7:** Average PD-1/PD-L1 expression on the mixed tumor environment. The bolded cells indicate which cells are being measured.

| Experiment | Microenvironments | Patient A | Patient B |
| --- | --- | --- | --- |
| PD-1 (**T cell**) | T cell / Tumor / Macrophages | 0.01886 | 0.00131 |
|  | T cell / Endothelial / Macrophages | 0.00558 | 0.00107 |
| PD-L1 (**Tumor**) | T cell / Tumor / Macrophages | **0.00649** | **0.00100** |
|  | Endothelial / Tumor / Macrophages | 0.00279 | 0.00046 |
| PD-L1 (**Macrophages**) | T cell / Tumor / Macrophages | **0.00204** | **0.00057** |
|  | T cell / Endothelial / Macrophages | 0.00068 | 0.00047 |

**Table 8:**
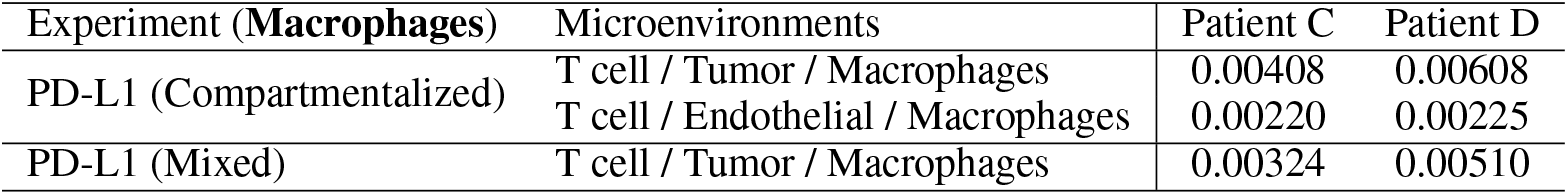
Average PDL1 expression of macrophages/monocytes on the compartmentalized tumor environment.

| Experiment (**Macrophages**) | Microenvironments | Patient C | Patient D |
| --- | --- | --- | --- |
| PD-L1 (Compartmentalized) | T cell / Tumor / Macrophages | 0.00408 | 0.00608 |
|  | T cell / Endothelial / Macrophages | 0.00220 | 0.00225 |
| PD-L1 (Mixed) | T cell / Tumor / Macrophages | 0.00324 | 0.00510 |

#### 3.6 T-CyCIF PD-1/PD-L1 Trend Experiment (Fig. 4C Left) Explanations

By training CCIGAN models on different segments of the t-CyCIF dataset (S2.2.2), we investigate the relationship between PD-1 and PD-L1. Using the same experimental setting as S3.4, our cell of interest (center cell) is a CD8 T cell and we iteratively add tumor cells as adjacent neighboring cells. Fig. 24 shows an example patch of a CD8 T cell (orange) in the center and iteratively adding tumor cells (yellow). As surrounding tumor cell surface area increases and surrounding PD-L1 expression increases, we expect PD-1 in the CD8 T cell to be upregulated, as it is an indicator of T cell exhaustion. PD-1 expression trend differs depending on the level of tumor-infiltration in the tumor microenvironment. In a poorly infiltrated microenvironment, the PD-1 expression trend should be greater than in a highly infiltrated microenvironment, since low infiltration indicates greater immunosuppression and a higher rate of T cell exhausation. Our results as shown in main paper Fig. 4C Left are fully displayed in Fig. 20, which illustrates sample runs, and table 9, which shows the full trend and statistical values from Fig. 4C Left. Additionally, we plotted the slope trend of CD8 T cell PD-1 expression against surrounding total tumor PD-L1 expression in table 10

**Table 9:** Slope and statistical values for t-CyCIF PD-1/PD-L1 Trend Experiment (S3.7) with respect to surrounding tumor cell surface area.

| Environment | $T_1$ | $T_2$ | $T_3$ |
| --- | --- | --- | --- |
| Slope | $2.629 \times 10^{-3}$ | $2.953 \times 10^{-3}$ | $3.134 \times 10^{-3}$ |
| $t$-test | 36.715 | 14.341 | 23.803 |
| $p$-value | $3.079 \times 10^{-225}$ | $1.945 \times 10^{-44}$ | $2.848 \times 10^{-110}$ |

**Table 10:** Slope and statistical values for t-CyCIF PD-1/PD-L1 Trend Experiment with respect to total surrounding PD-L1 expression.

| Environment | $T_1$ | $T_2$ | $T_3$ |
| --- | --- | --- | --- |
| Slope | $4.339 \times 10^{-5}$ | $5.450 \times 10^{-5}$ | $2.970 \times 10^{-4}$ |
| $t$-test | 35.248 | 14.477 | 25.608 |
| $p$-value | $2.098 \times 10^{-211}$ | $3.285 \times 10^{-45}$ | $4.520 \times 10^{-125}$ |

**Figure 20:**
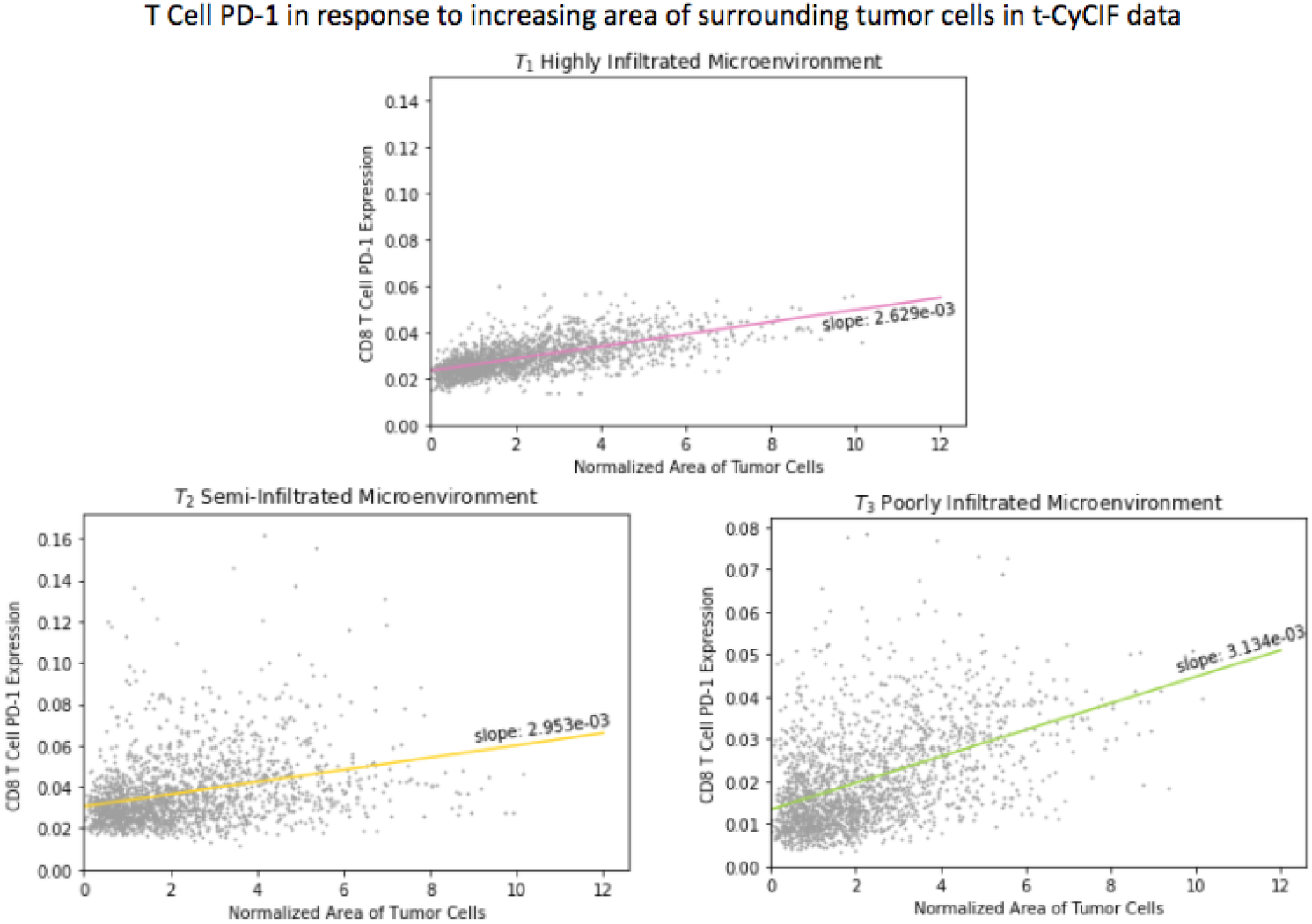
PD-1/PD-L1 experiment on tumor microenvironments with varying levels of tumor-infiltration in t-CyCIF (S3.6) with respect to surrounding tumor cell surface area.

**Figure 21:**
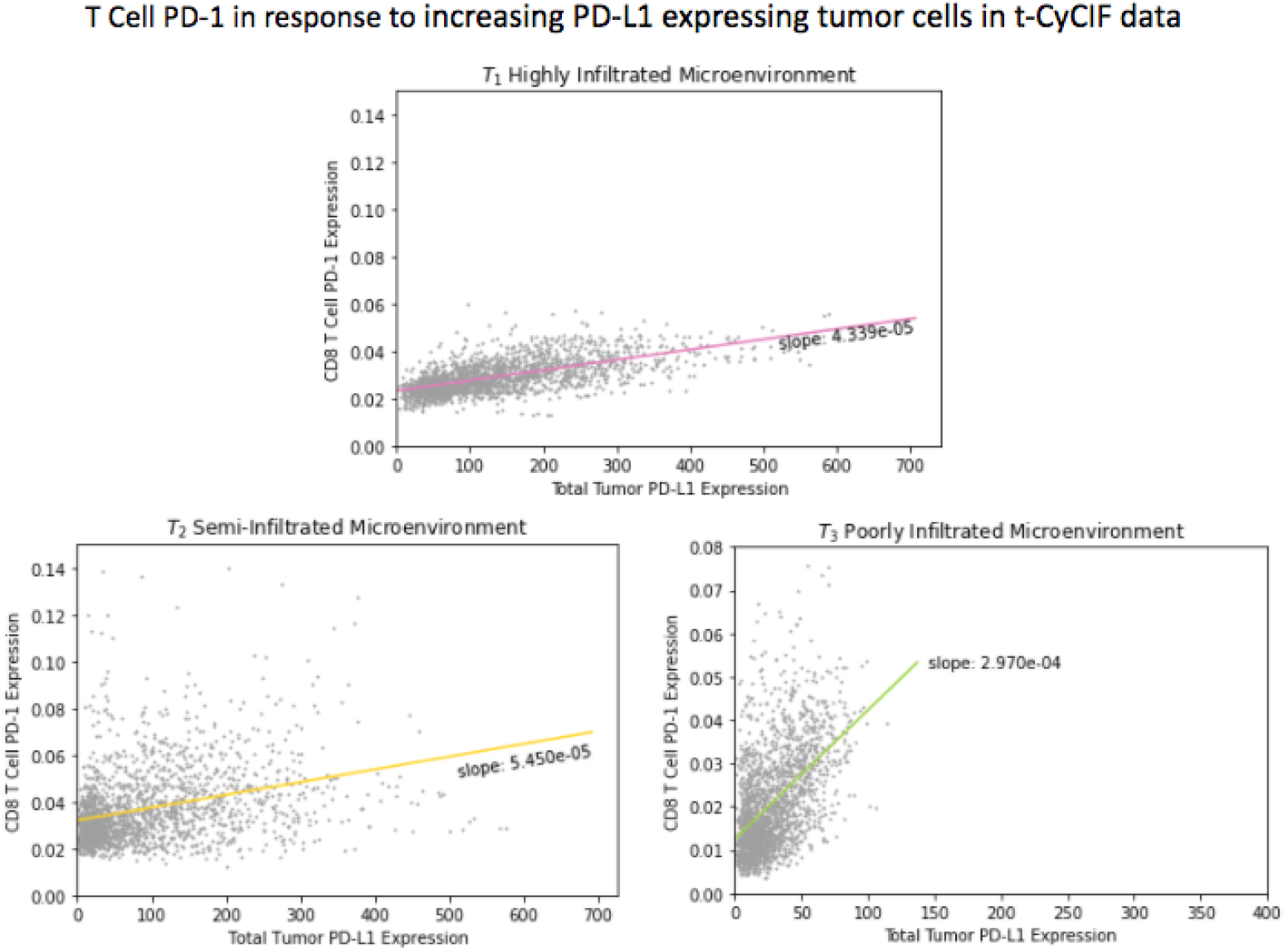
PD-1/PD-L1 experiment on tumor microenvironments with varying levels of tumor-infiltration in t-CyCIF (S3.6) with respect to total surrounding PD-L1 expression.

**Table 11:** Slope and statistical values for t-CyCIF LAG3/PD-L1 Trend Experiment.

| Environment | $T_1$ | $T_2$ | $T_3$ |
| --- | --- | --- | --- |
| Slope | $4.843 \times 10^{-5}$ | $8.838 \times 10^{-5}$ | $6.242 \times 10^{-4}$ |
| $t$-test | 35.778 | 12.711 | 21.693 |
| $p$-value | $2.172 \times 10^{-216}$ | $1.238 \times 10^{-35}$ | $9.723 \times 10^{-94}$ |

#### 3.7 T-CyCIF LAG3/PD-L1 Trend Experiment (Fig. 4C Right) Explanations

Furthermore, we investigate the relationship between LAG3 and PD-L1. Using the same experimental setting as S3.6, our cell of interest (center cell) is a CD8 T cell and the adjacent neighboring cells are tumor cells. As tumor cells are added and the surrounding PD-L1 expression increases, we expect LAG3 in the CD8 T cell to be upregulated, as it is another indicator of T cell exhaustion. Similarly to S3.6, we expect a higher trend of LAG3 upregulation with respect to PD-L1 in microenvironments with lower tumor-infiltration. Our results as shown in main paper Fig. 4C Right are fully displayed in Figure 22, which illustrates sample runs, and table 11, which shows the full trend and statistical values from Fig. 4C Right.

**Figure 22:**
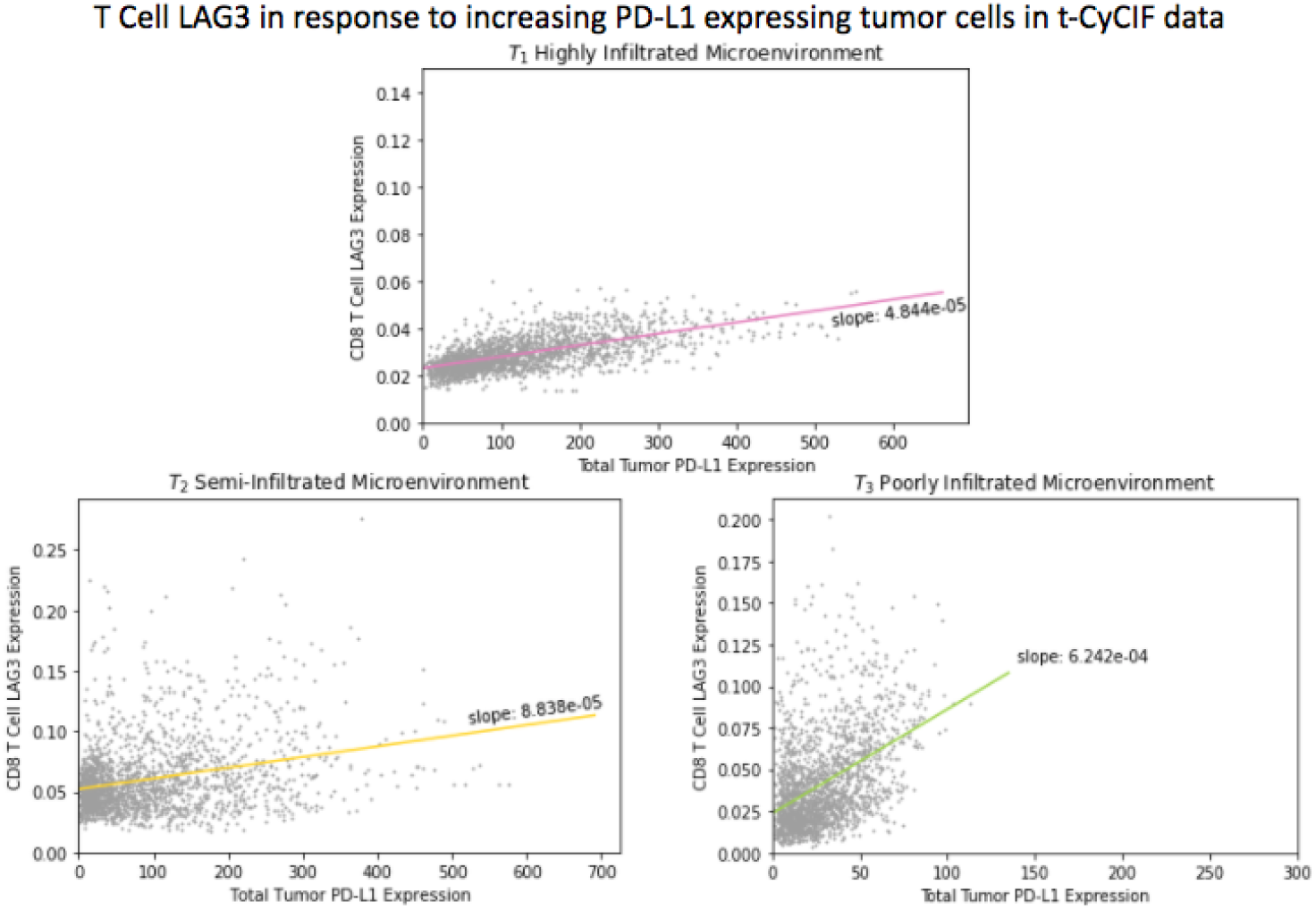
LAG3/PD-1 experiment on tumor microenvironments with varying levels of tumor-infiltration in t-CyCIF

### 4 Further Experiments and Experiment Details

#### 4.1 Experimental Setup

In our general experiments that involved iteratively manipulating cell patches, we created a experimental dataset of approximately 1000 patches for each type of data (MIBI-TOF or t-CyCIF) and manipulated the cell types to the necessary cell types for each experiment. For a patch *s* in the experimental dataset with *n* cells in the patch, we expanded the patch into *n* individual patches 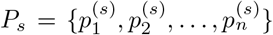 where for 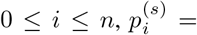 patch of cells from 1 to *i*. An example can be seen in Fig. 23 for MIBI data where PD-1 expression in the CD 8 T cell of interest reacts to newly introduced PD-L1 expressing tumor cells. Another example is shown for t-CyCIF data in Fig. 24. Following this, changes in protein expressions in the cell of interest due to newly introduced cells were analyzed using a variety of techniques (center of mass, summation, mass shift, regression trend).

**Figure 23:**
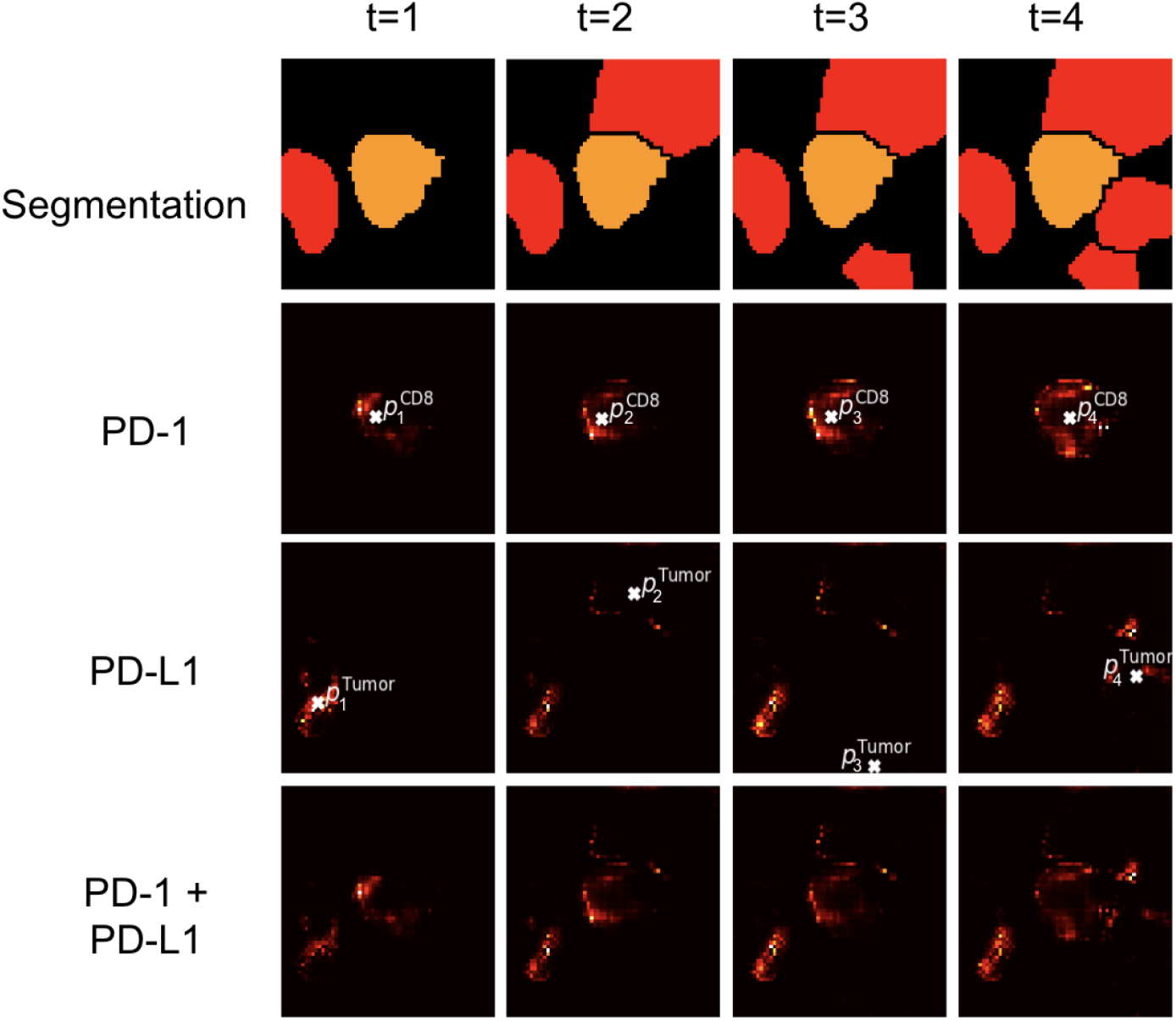
Process of iteratively adding tumor cells in MIBI-TOF. The added red cells are tumor cells (PD-L1) and the center orange cell indicates a CD8 T cell (cell of interest, PD-1). For this process, we focus on each instance of an added tumor.

**Figure 24:**
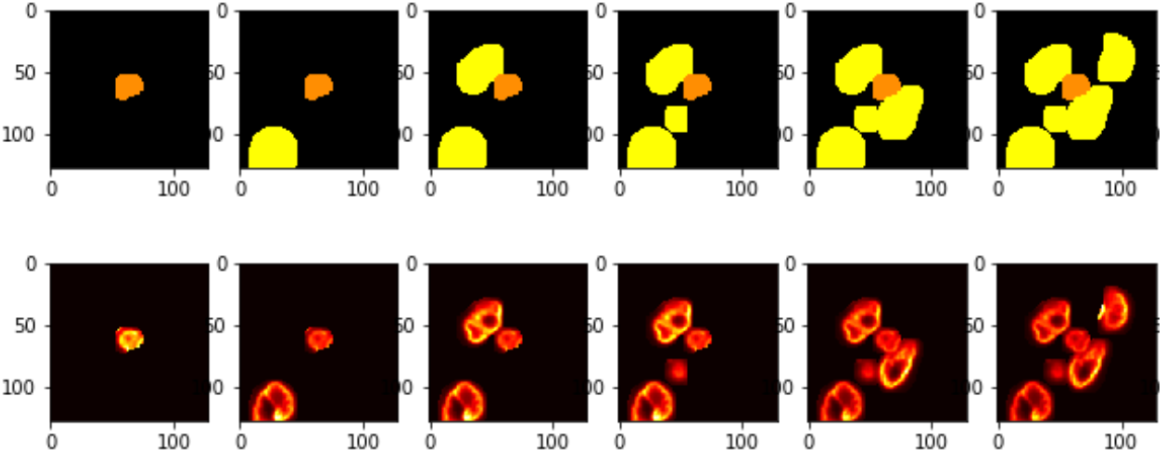
Iteratively Adding Cells for t-CyCIF data. A random dsDNA protein channel is shown.

#### 4.2 Search Algorithm Discussion

In this section we discuss potential biological trends, erroneous correlations, and algorithm settings. It is important to note that some of the spurious correlations are easily explained by poor segmentations (known issue).

Fig. 5 displays the results of our search algorithm, which shows which particular cell-cell interactions are significantly captured in CCIGAN. For four primary cells– tumor, macrophage, CD8, and endothelial, we measure their expression level change in a subset of protein markers and calculate the relative frequency of this cell-cell interaction. Using a subset of 1268 test cell patches, we counted the number of significant logged interactions that produced an increase or decrease in expression level greater than a chosen threshold. Expression level change was measured by simply calculating the difference in summed pixel intensity (protein expression) in a particular channel.

For the relative frequency of a particular cell-cell interaction, the normalization schema is as follows. Using the logged counts of significant changes in expression level, we take the max of either the increase counts or the decrease counts. Then, to quantify the ratio of relative importance within a primary cell group, we divide the max counts by the total number of logged counts (for either only increases or decreases). For example, say we are measuring tumor as the primary cell. If 50 decreases and 200 increases in expression level are recorded, with 1000 total increases logged for the tumor primary cell, 200*/*1000 = 0.2 is the relative frequency after normalization.

Algorithm settings were set on the same sensitivity threshold for experiments. Additionally, figure generation normalization was done across protein markers (within different cell groups). Finally, interactions comprising of 5% or less of total logged interactions were disregarded.

Individual marker discussions are given below:

1. Vimentin It was observed that nearly all scenarios of cell-cell interaction accounted for by CCIGAN resulted in an varying increases of vimentin expression in the cell of interest. While not necessarily biologically explained, the observed changes are plausible as vimentin is a structural protein found in all cells and changes in its levels may not be attributable to specific cell interactions. A continued investigation is given in S4.3 regarding tumor and CD 8 T cell vimentin expressions.
2. PD-1 A slight increase in the PD-1 expression is seen on tumor cells when surrounded by CD8 T cells. However, this increase was negligible and can be attributed to noisy segmentations where PD-1 expressing CD8 T cells are located near tumor cells, suggesting in the training data that a tumor cell expresses PD-1.
3. PD-L1 The model suggests PD-L1 expression on CD8 T cells. This is due to noisy segmentations similar to the situation in PD-1.
4. CD8 The model predicted an increase in CD8 expression on the tumor cell in several scenarios despite this not being biologically expected. This also due to the same noisy segmentations.
5. Pan Keratin The decreased levels in Pan Keratin expression within the tumor cell line with neighboring non-tumor cells was biologically expected. However, the slight increase in pan keratin expression when the tumor cell of interest had tumor cell neighbors may be due to the same noisy segmentations as mentioned before.

#### 4.3 Vimentin Table Expressions

We measure trends of vimentin expression in tumor cells due to varying amounts of surrounding CD 8 T cells. The cell of interest is a tumor cell and we iteratively change surrounding adjacent cells from tumor cells to CD8 T cells with different probabilities. Then we measure the number of instances a change of vimentin is detected, and what that change is.

Vimentin plays an important role in tumorigenesis both as a structural protein and in cellular signalling. Numerous studies have noted vimentin’s regulatory importance in a variety of cell signalling pathways which promote tumor survival and resistance to cellular stress (41) (42). CCIGAN experiments wherein a tumor cell of interest was surrounded by increasing numbers of CD8 T cells caused up to a 91.6% increase in the total tumor vimentin expression (S4.2.1). Increasing CD8 T cell presence around a tumor cell is likely to be associated with increasing amounts of pro-apoptotic signalling on the tumor cell, causing a large degree of cellular stress. Thus the increase in the tumor’s vimentin would be a biologically expected outcome given the protein’s role in promoting cellular survival pathways.

**Table 12:** Table indicating net changes in Vimentin expression in a tumor cells due to increasing CD8 T cell presence.

| Cell of Interest | Adjacent Cell Neighbor % | % of Patches Indicating Increase in Vimentin |
| --- | --- | --- |
| Tumor | 100% CD8 0% Tumor | 91.6% |
| Tumor | 75% CD8 25% Tumor | 81.6% |
| Tumor | 50% CD8 50% Tumor | 60.4% |
| Tumor | 25% CD8 75% Tumor | 26.7% |

**Table 13:** CCIGAN Biological Discovery Evaluation

| Patient | CD8/PK Exp. Slope | CD8/PK Cont. Slope | EM (exp.) | EM control |
| --- | --- | --- | --- | --- |
| 3 | -4.36E-06 | -2.04E-06 | 8.98 | 3.31 |
| 4 | -8.99E-09 | 3.15E-07 | 33.66 | 21.49 |
| 5 | -3.05E-06 | 3.01E-07 | 21.77 | 5.97 |
| 9 | -2.66E-06 | -9.11E-07 | 4.99 | -0.35 |
| 10 | 2.41E-06 | 1.39E-06 | 74.34 | 41.55 |
| 13 | -4.86E-06 | 3.70E-06 | 79.84 | -6.64 |
| 14 | -2.87E-06 | 1.52E-07 | 151.56 | -22.68 |
| 16 | -2.13E-05 | 2.66E-204 | 12.53 | -4.33 |
| 17 | -2.71E-07 | 4.98E-06 | 37.01781478 | -44.33682343 |
| 19 | -1.36E-07 | 4.18E-07 | 39.09 | -35.229 |
| 20 | -1.50E-07 | 2.62E-07 | 4.21 | 1.05 |
| 29 | -2.60E-07 | 3.84E-06 | 4.89 | 2.74 |
| 35 | -1.10E-07 | 1.90E-05 | 4.63 | 2.96 |
| 37 | -1.72E-05 | -7.50E-06 | 32.01 | 6.42 |
| 39 | -6.18E-07 | 1.77E-06 | 5.09 | 0.13 |

**Table 14:** SPADE Biological Discovery Evaluation

| Patient | CD8/PK Exp. Slope | CD8/PK Cont. Slope | EM (exp.) | EM control |
| --- | --- | --- | --- | --- |
| 3 | -4.57E-06 | -2.70E-06 | -2.89 | -9.49 |
| 4 | 5.45E-07 | 6.48E-07 | 24.3 | -4.73 |
| 5 | -3.97E-06 | -1.23E-06 | -0.96 | -6.52 |
| 9 | -4.54E-06 | -2.54E-06 | -35.2 | -64.64 |
| 10 | -1.07E-06 | -1.04E-06 | -33.3 | -43.26 |
| 13 | -1.47E-06 | 6.57E-06 | 5.8 | -20.2 |
| 14 | -1.69E-06 | 1.64E-06 | 13.13 | -61.91 |
| 16 | -9.71E-06 | -8.27E-06 | -0.07 | -0.082 |
| 17 | -6.57E-06 | -3.08E-07 | -4.94 | -6.729 |
| 19 | 2.37E-06 | 1.14E-06 | -0.15 | 0.425 |
| 20 | -1.18E-06 | -9.33E-07 | -10.77 | -2.77 |
| 29 | -3.84E-07 | 3.78E-06 | -0.68 | 2.43 |
| 35 | 2.20E-05 | 3.60E-05 | 6.97 | 2.669 |
| 37 | -1.35E-05 | -6.39E-06 | -3.75 | 3.846 |
| 39 | -1.40E-05 | -8.07E-06 | -0.47 | -3.3 |

**Table 15:**
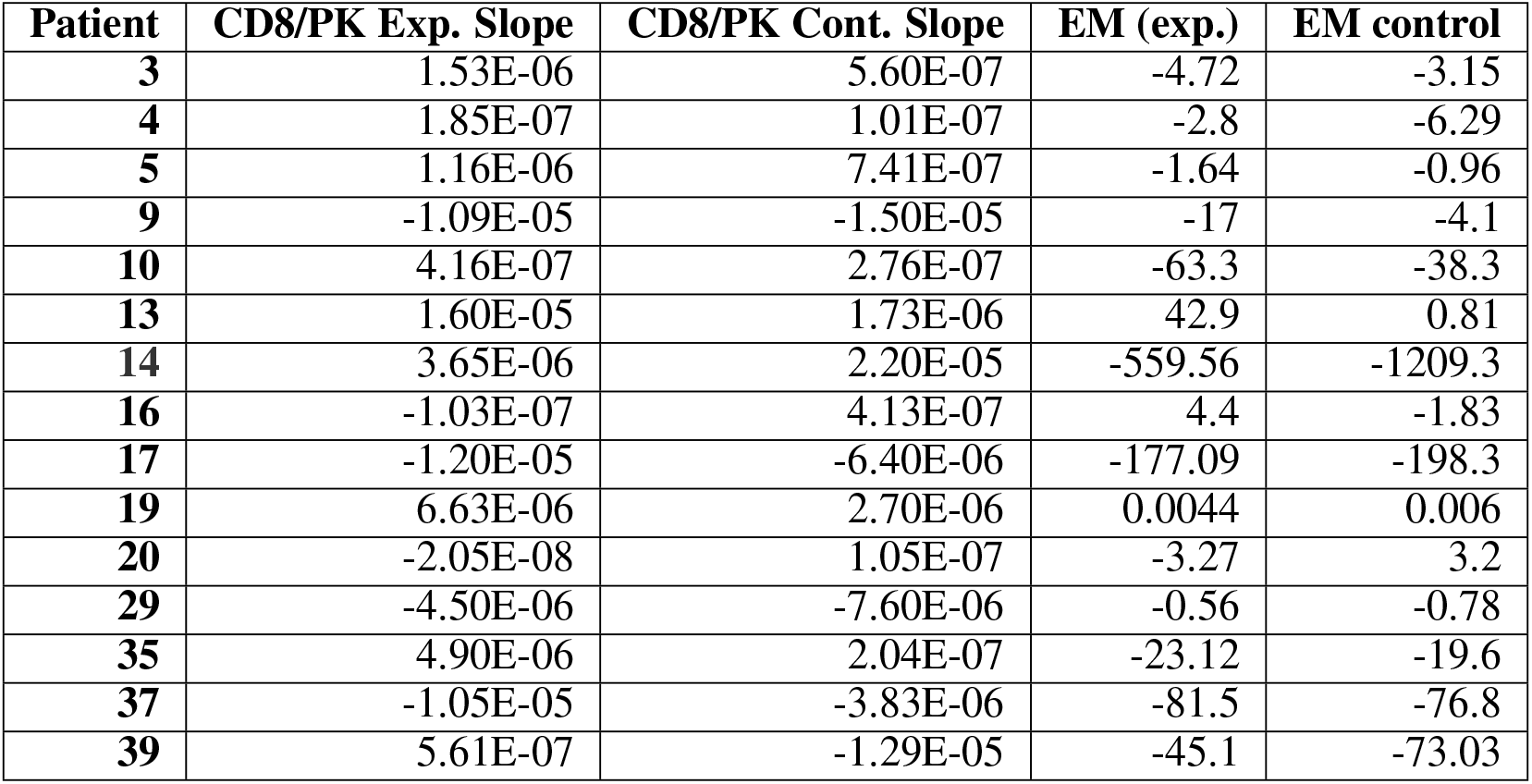
Pix2Pix Biological Discovery Evaluation

**Table 16:**
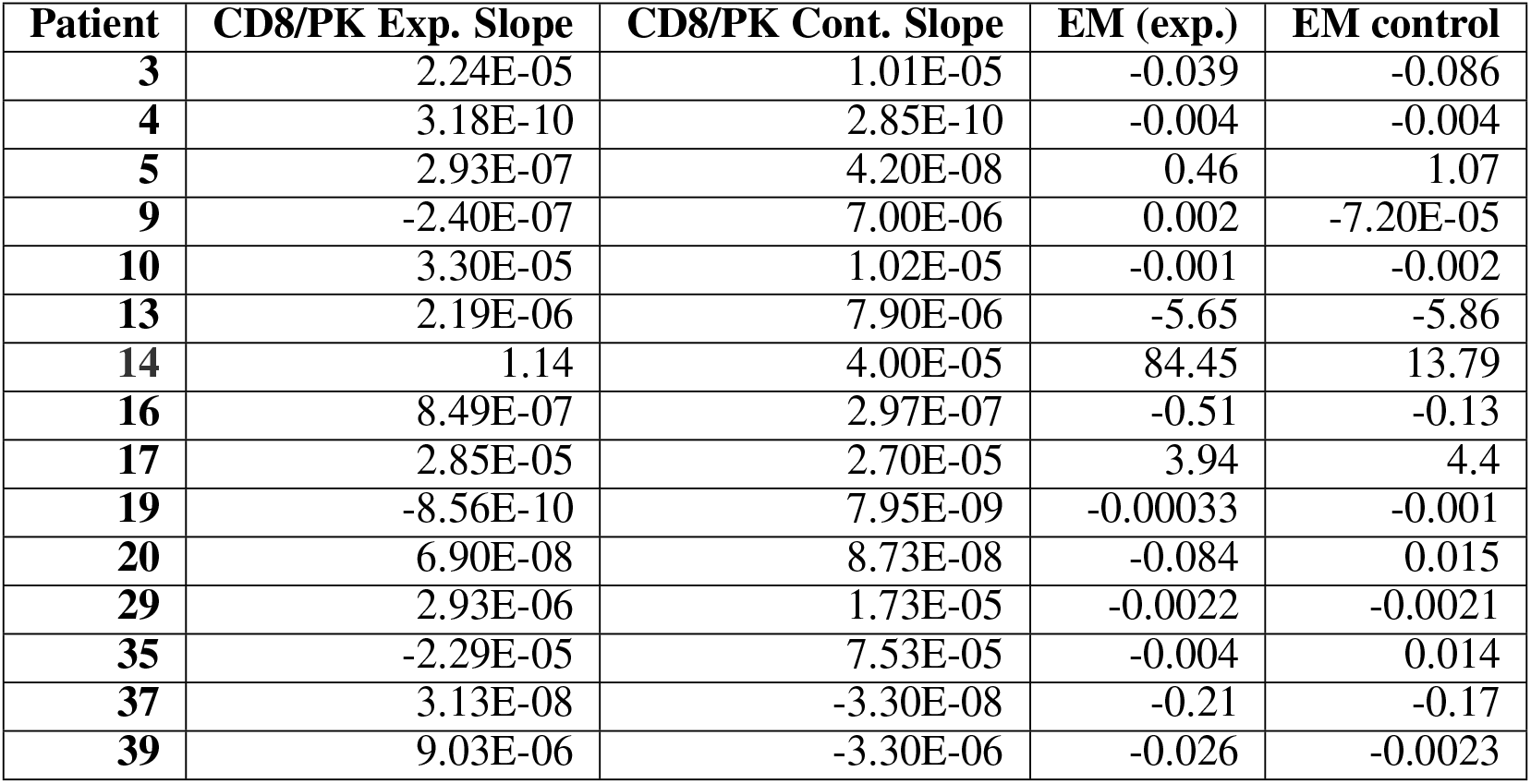
CycleGAN Biological Discovery Evaluation

| Patient | CD8/PK Exp. Slope | CD8/PK Cont. Slope | EM (exp.) | EM control |
| --- | --- | --- | --- | --- |
| 3 | 2.24E-05 | 1.01E-05 | -0.039 | -0.086 |
| 4 | 3.18E-10 | 2.85E-10 | -0.004 | -0.004 |
| 5 | 2.93E-07 | 4.20E-08 | 0.46 | 1.07 |
| 9 | -2.40E-07 | 7.00E-06 | 0.002 | -7.20E-05 |
| 10 | 3.30E-05 | 1.02E-05 | -0.001 | -0.002 |
| 13 | 2.19E-06 | 7.90E-06 | -5.65 | -5.86 |
| 14 | 1.14 | 4.00E-05 | 84.45 | 13.79 |
| 16 | 8.49E-07 | 2.97E-07 | -0.51 | -0.13 |
| 17 | 2.85E-05 | 2.70E-05 | 3.94 | 4.4 |
| 19 | -8.56E-10 | 7.95E-09 | -0.00033 | -0.001 |
| 20 | 6.90E-08 | 8.73E-08 | -0.084 | 0.015 |
| 29 | 2.93E-06 | 1.73E-05 | -0.0022 | -0.0021 |
| 35 | -2.29E-05 | 7.53E-05 | -0.004 | 0.014 |
| 37 | 3.13E-08 | -3.30E-08 | -0.21 | -0.17 |
| 39 | 9.03E-06 | -3.30E-06 | -0.026 | -0.0023 |

**Table 17:** CCIGAN Reconstruction Metrics

| Patient | COM (Exp.) | Random COM | SSIM | Adjust L1 | Pure L1 | Adjust MSE | Pure MSE | Cell MI |
| --- | --- | --- | --- | --- | --- | --- | --- | --- |
| 3 | 10.45 | 12.13 | 0.78 | 1.4034 | 1.353 | 0.07329 | 0.07312 | 4.315 |
| 4 | 10.04 | 12.39 | 0.73 | 1.502 | 1.4759 | 0.1142 | 0.11399 | 1.18 |
| 5 | 9.67 | 11.77 | 0.81 | 1.4017 | 1.3003 | 0.086759 | 0.08644 | 4.93 |
| 9 | 10.22 | 11.78 | 0.82 | 1.0385 | 0.98948 | 0.0527 | 0.05257 | 7.63 |
| 10 | 9.98 | 11.88 | 0.81 | 0.93256 | 0.89 | 0.0462 | 0.0458 | 9.15 |
| 13 | 10.39 | 11.97 | 0.77 | 1.76 | 1.69 | 0.086 | 0.0855 | 36.8 |
| 14 | 10.46 | 12.27 | 0.81 | 0.613 | 0.594 | 0.026 | 0.026 | 10.46 |
| 16 | 10.72 | 12.53 | 0.72 | 1.992 | 1.945 | 0.114 | 0.114 | 10.83 |
| 17 | 10.56 | 12.16 | 0.71 | 2.14 | 2.008 | 0.109 | 0.107 | 16.16 |
| 19 | 10.71 | 12.38 | 0.9 | 0.308 | 0.299 | 0.0194 | 0.0194 | 0.044 |
| 20 | 10.469 | 11.93 | 0.77 | 1.093 | 1.0119 | 0.055 | 0.054 | 14.14 |
| 29 | 10.03 | 12.443 | 0.8 | 1.057 | 1.031 | 0.075 | 0.075 | 10.08 |
| 35 | 10.14 | 11.66 | 0.83 | 1.225 | 1.175 | 0.065 | 0.065 | 1.53 |
| 37 | 10.44 | 12.57 | 0.84 | 0.947 | 0.916 | 0.056 | 0.0557 | 6.84 |
| 39 | 9.65 | 11.83 | 0.84 | 1.24 | 1.01 | 0.05 | 0.049 | 18.38 |

**Table 18:** SPADE Reconstruction Metrics

| Patient | COM (Exp.) | Random COM | SSIM | Adjust L1 | Pure L1 | Adjust MSE | Pure MSE | Cell MI |
| --- | --- | --- | --- | --- | --- | --- | --- | --- |
| 3 | 11.12 | 12.32 | 0.763 | 1.26 | 1.23 | 0.072 | 0.072 | 3.933 |
| 4 | 10.61 | 12.24 | 0.734 | 1.52 | 1.5 | 0.12 | 0.12 | 1.264 |
| 5 | 10.5 | 11.93 | 0.8 | 1.05 | 1.04 | 0.088 | 0.088 | 3.843 |
| 9 | 11.26 | 12.1 | 0.8 | 0.89 | 0.87 | 0.056 | 0.056 | 6.474 |
| 10 | 10.98 | 12.04 | 0.78 | 0.863 | 0.84 | 0.049 | 0.049 | 8.002 |
| 13 | 11.3 | 12.09 | 0.74 | 1.66 | 1.64 | 0.096 | 0.096 | 32.78 |
| 14 | 10.8 | 11.99 | 0.77 | 0.89 | 0.87 | 0.055 | 0.055 | 9.99 |
| 16 | 11.47 | 12.68 | 0.74 | 1.79 | 1.77 | 0.096 | 0.093 | 10.98 |
| 17 | 11.435 | 12.24 | 0.73 | 1.64 | 1.6 | 0.1 | 0.1 | 13.215 |
| 19 | 11.05 | 12.63 | 0.89 | 0.286 | 0.281 | 0.02 | 0.02 | 0.037 |
| 20 | 11.24 | 12.15 | 0.73 | 0.96 | 0.94 | 0.063 | 0.063 | 12.26 |
| 29 | 11.07 | 12.52 | 0.81 | 1.11 | 1.104 | 0.071 | 0.071 | 11.586 |
| 35 | 10.48 | 12.16 | 0.79 | 1.22 | 1.2 | 0.078 | 0.077 | 1.299 |
| 37 | 11.6 | 12.52 | 0.834 | 0.82 | 0.81 | 0.051 | 0.051 | 6.301 |
| 39 | 10.58 | 11.91 | 0.8 | 0.9 | 0.88 | 0.05 | 0.05 | 15.618 |

**Table 19:** Pix2Pix Reconstruction Metrics

| Patient | COM (Exp.) | Random COM | SSIM | Adjust L1 | Pure L1 | Adjust MSE | Pure MSE | Cell MI |
| --- | --- | --- | --- | --- | --- | --- | --- | --- |
| 3 | 13.05 | 12.85 | 0.64 | 1.76 | 1.58 | 0.13 | 0.12 | 3.153 |
| 4 | 12.06 | 12.15 | 0.6 | 2.11 | 1.9 | 0.18 | 0.17 | 1.014 |
| 5 | 12.6 | 12.4 | 0.67 | 1.26 | 1.14 | 0.12 | 0.12 | 2.884 |
| 9 | 11.89 | 12.11 | 0.71 | 1.06 | 0.95 | 0.07 | 0.066 | 5.12 |
| 10 | 11 | 12.53 | 0.65 | 1.29 | 1.16 | 0.16 | 0.14 | 6.29 |
| 13 | 10.54 | 12.9 | 0.58 | 2.75 | 2.48 | 0.24 | 0.23 | 30.405 |
| 14 | 11.55 | 12.5 | 0.63 | 1.22 | 1.09 | 0.088 | 0.082 | 9.708 |
| 16 | 10.5 | 12.2 | 0.67 | 2.26 | 2.07 | 0.12 | 0.11 | 10.833 |
| 17 | 11.15 | 12.46 | 0.59 | 2.32 | 2.04 | 0.07 | 0.155 | 11.703 |
| 19 | 11.74 | 11.94 | 0.84 | 0.46 | 0.38 | 0.037 | 0.033 | 0.027 |
| 20 | 10.88 | 12.02 | 0.66 | 1.15 | 1.07 | 0.085 | 0.082 | 11.64 |
| 29 | 11.6 | 12.66 | 0.7 | 1.43 | 1.28 | 0.12 | 0.11 | 9.49 |
| 35 | 11.98 | 12.18 | 0.67 | 1.56 | 1.34 | 0.11 | 0.1 | 1.017 |
| 37 | 11.2 | 11.9 | 0.75 | 1.19 | 1.05 | 0.08 | 0.07 | 5.478 |
| 39 | 11.07 | 12.51 | 0.74 | 1.09 | 0.99 | 0.097 | 0.09 | 12.253 |

**Table 20:** CycleGAN Reconstruction Metrics

| Patient | COM (Exp.) | Random COM | SSIM | Adjust L1 | Pure L1 | Adjust MSE | Pure MSE | Cell MI |
| --- | --- | --- | --- | --- | --- | --- | --- | --- |
| 3 | 13.7 | 12.8 | 0.386 | 6.26 | 5.214 | 2.46 | 2.02 | 2.812 |
| 4 | 13.01 | 12.49 | 0.406 | 7.924 | 6.43 | 3.654 | 2.946 | 0.888 |
| 5 | 9.96 | 12.49 | 0.5 | 2.73 | 2.44 | 0.712 | 0.656 | 2.51 |
| 9 | 12.295 | 12.56 | 0.49 | 2.51 | 2.18 | 0.64 | 0.555 | 3.904 |
| 10 | 15.27 | 12.34 | 0.35 | 5.99 | 4.89 | 2.88 | 2.42 | 5.722 |
| 13 | 11.21 | 12.34 | 0.29 | 9.42 | 8.1 | 4.35 | 3.73 | 21.91 |
| 14 | 13.29 | 12.29 | 0.31 | 6.66 | 5.4 | 2.63 | 2.11 | 8.23 |
| 16 | 13.97 | 12.2 | 0.25 | 8.75 | 7.5 | 3.1 | 2.65 | 6.735 |
| 17 | 12.74 | 12.6 | 0.29 | 11.48 | 9.6 | 5.13 | 4.3 | 9.019 |
| 19 | 10.97 | 12.27 | 0.83 | 0.6 | 0.53 | 0.129 | 0.1146 | 0.021 |
| 20 | 13.95 | 12.5 | 0.3 | 7.24 | 5.92 | 3.46 | 2.83 | 9.736 |
| 29 | 13.36 | 12.59 | 0.505 | 3.97 | 3.55 | 1.165 | 1.062 | 8.668 |
| 35 | 11.67 | 12.27 | 0.41 | 7.24 | 5.75 | 3.03 | 2.25 | 0.982 |
| 37 | 13.2 | 12.7 | 0.5 | 4.52 | 4 | 1.74 | 1.6 | 3.967 |
| 39 | 11.68 | 12.2 | 0.4 | 3.69 | 3.11 | 1.14 | 0.97 | 10.7 |

## Footnotes

1 These 4 patients were chosen because they best demonstrate clearly delineated compartmentalized and mixed TMEs.

## References

[1] Gregory L. Beatty and Whitney L. Gladney. Immune escape mechanisms as a guide for cancer immunotherapy. Clinical Cancer Research, 21(4):687–692, 2015. ISSN 1078-0432. doi: 10.1158/1078-0432.CCR-14-1860. URL https://clincancerres.aacrjournals.org/content/21/4/687.

[2] Els M. E. Verdegaal, Noel F. C. C. de Miranda, Marten Visser, Tom Harryvan, Marit M. van Buuren, Rikke S. Andersen, Sine R. Hadrup, Caroline E. van der Minne, Remko Schotte, Hergen Spits, John B. A. G. Haanen, Ellen H. W. Kapiteijn, Ton N. Schumacher, and Sjoerd H. van der Burg. Neoantigen landscape dynamics during human melanoma–t cell interactions. Nature, 536(7614):91–95, 2016. doi: 10.1038/nature18945. URL https://doi.org/10.1038/nature18945.

[3] Christian M Schürch, Salil S Bhate, Graham L Barlow, Darci J Phillips, Luca Noti, Inti Zlobec, Pauline Chu, Sarah Black, Janos Demeter, David R McIlwain, et al. Coordinated cellular neighborhoods orchestrate antitumoral immunity at the colorectal cancer invasive front. Cell, 182(5):1341–1359, 2020.

[4] Jia-Ren Lin, Benjamin Izar, Shu Wang, Clarence Yapp, Shaolin Mei, Parin M Shah, Sandro Santagata, and Peter K Sorger. Highly multiplexed immunofluorescence imaging of human tissues and tumors using t-cycif and conventional optical microscopes. Elife, 7, 2018.

[5] Leeat Keren, Marc Bosse, Steve Thompson, Tyler Risom, Kausalia Vijayaragavan, Erin McCaffrey, Diana Marquez, Roshan Angoshtari, Noah F Greenwald, Harris Fienberg, et al. Mibitof: A multiplexed imaging platform relates cellular phenotypes and tissue structure. Science advances, 5(10):eaax5851, 2019.

[6] Michael Angelo, Sean C Bendall, Rachel Finck, Matthew B Hale, Chuck Hitzman, Alexander D Borowsky, Richard M Levenson, John B Lowe, Scot D Liu, Shuchun Zhao, Yasodha Natkunam, and Garry P Nolan. Multiplexed ion beam imaging of human breast tumors. Nature medicine, 20(4):436–442, 04 2014. doi: 10.1038/nm.3488. URL https://pubmed.ncbi.nlm.nih.gov/24584119.

[7] Leeat Keren, Marc Bossé, Diana Marquez, Roshan Angoshtari, Samir Jain, Sushama Varma, Soo-Ryum Yang, Allison Kurian, David Valen, Robert West, Sean Bendall, and Michael Angelo. A structured tumor-immune microenvironment in triple negative breast cancer revealed by multiplexed ion beam imaging. Cell, 174:1373–1387.e19, 09 2018. doi: 10.1016/j.cell.2018.08.039.

[8] Hartland W. Jackson, Jana R. Fischer, Vito R. T. Zanotelli, H. Raza Ali, Robert Mechera, Savas D. Soysal, Holger Moch, Simone Muenst, Zsuzsanna Varga, Walter P. Weber, and Bernd Bodenmiller. The single-cell pathology landscape of breast cancer. Nature, 578(7796):615–620, 2020. doi: 10.1038/s41586-019-1876-x. URL https://doi.org/10.1038/s41586-019-1876-x.

[9] Damien Arnol, Denis Schapiro, Bernd Bodenmiller, Julio Saez-Rodriguez, and Oliver Stegle. Modeling cell-cell interactions from spatial molecular data with spatial variance component analysis. Cell Reports, 29(1):202 – 211.e6, 2019. ISSN 2211-1247. doi: https://doi.org/10.1016/j.celrep.2019.08.077. URL http://www.sciencedirect.com/science/article/pii/S2211124719311325.

[10] Piotr Baniukiewicz, E. Josiah Lutton, Sharon Collier, and Till Bretschneider. Generative adversarial networks for augmenting training data of microscopic cell images. Frontiers in Computer Science, 1:10, 2019. ISSN 2624-9898. doi: 10.3389/fcomp.2019.00010. URL https://www.frontiersin.org/article/10.3389/fcomp.2019.00010.

[11] Anton Osokin, Anatole Chessel, Rafael Edgardo Carazo-Salas, and Federico Vaggi. Gans for biological image synthesis. CoRR, abs/1708.04692, 2017. URL http://arxiv.org/abs/1708.04692.

[12] Gregory R. Johnson, Rory M. Donovan-Maiye, and Mary M. Maleckar. Generative modeling with conditional autoencoders: Building an integrated cell, 2017.

[13] Ting Zhao and Robert F. Murphy. Automated learning of generative models for subcellular location: Building blocks for systems biology. Cytometry Part A, 71A(12):978–990, 2007. doi: 10.1002/cyto.a.20487. URL https://onlinelibrary.wiley.com/doi/abs/10.1002/cyto.a.20487.

[14] Taesung Park, Ming-Yu Liu, Ting-Chun Wang, and Jun-Yan Zhu. Semantic image synthesis with spatially-adaptive normalization. CoRR, abs/1903.07291, 2019. URL http://arxiv.org/abs/1903.07291.

[15] Norio Azumi and Hector Battifora. The distribution of vimentin and keratin in epithelial and nonepithelial neoplasms: a comprehensive immunohistochemical formalin-and alcohol-fixed tumors. American journal of clinical pathology, 88(3):286–296, 1987.

[16] Luis Martínez-Lostao, Alberto Anel, and Julián Pardo. How do cytotoxic lymphocytes kill cancer cells?, 2015.

[17] RG Oshima. Apoptosis and keratin intermediate filaments. Cell Death & Differentiation, 9(5):486–492, 2002.

[18] V Badock, U Steinhusen, K Bommert, B Wittmann-Liebold, and A Otto. Apoptosis-induced cleavage of keratin 15 and keratin 17 in a human breast epithelial cell line. Cell Death & Differentiation, 8(3):308–315, 2001.

[19] Nam-On Ku, Pavel Strnad, Heike Bantel, and M Bishr Omary. Keratins: Biomarkers and modulators of apoptotic and necrotic cell death in the liver. Hepatology, 64(3):966–976, 2016.

[20] Jun-Yan Zhu, Taesung Park, Phillip Isola, and Alexei A Efros. Unpaired image-to-image translation using cycle-consistent adversarial networks. In Computer Vision (ICCV), 2017 IEEE International Conference on, 2017.

[21] Mojgan Ahmadzadeh, Laura A. Johnson, Bianca Heemskerk, John R. Wunderlich, Mark E. Dudley, Donald E. White, and Steven A. Rosenberg. Tumor antigen–specific CD8 T cells infiltrating the tumor express high levels of PD-1 and are functionally impaired. Blood, 114(8):1537–1544, 08 2009. ISSN 0006-4971. doi: 10.1182/blood-2008-12-195792. URL https://doi.org/10.1182/blood-2008-12-195792.

[22] Shuming Chen, George A. Crabill, Theresa S. Pritchard, Tracee L. McMiller, Ping Wei, Drew M. Pardoll, Fan Pan, and Suzanne L. Topalian. Mechanisms regulating pd-l1 expression on tumor and immune cells. Journal for ImmunoTherapy of Cancer, 7(1):305, 2019.

[23] David Escors, María Gato-Cañas, Miren Zuazo, Hugo Arasanz, María Jesus García-Granda, Ruth Vera, and Grazyna Kochan. The intracellular signalosome of pd-l1 in cancer cells. Signal transduction and targeted therapy, 3:26–26, 09 2018.

[24] Ting-Chun Wang, Ming-Yu Liu, Jun-Yan Zhu, Andrew Tao, Jan Kautz, and Bryan Catanzaro. High-resolution image synthesis and semantic manipulation with conditional gans. In Proceedings of the IEEE Conference on Computer Vision and Pattern Recognition, 2018.

[25] Michael J Eppihimer, Jason Gunn, Gordon J Freeman, Edward A Greenfield, Tetyana Chernova, Jamie Erickson, and John P Leonard. Expression and regulation of the pd-l1 immunoinhibitory molecule on microvascular endothelial cells. Microcirculation (New York, N.Y. : 1994), 9(2):133–145, 04 2002.

[26] Nancy Rodig, Timothy Ryan, JessicaA. Allen, Hong Pang, Nir Grabie, Tatyana Chernova, EdwardA. Greenfield, Spencer C. Liang, ArleneH. Sharpe, AndrewH. Lichtman, and GordonJ. Freeman. Endothelial expression of pd-l1 and pd-l2 down-regulates cd8+ t cell activation and cytolysis. European Journal of Immunology, 33(11):3117–3126, 2003. doi: 10.1002/eji.200324270. URL https://onlinelibrary.wiley.com/doi/abs/10.1002/eji.200324270.

[27] Rumana Rashid, Giorgio Gaglia, Yu-An Chen, Jia-Ren Lin, Ziming Du, Zoltan Maliga, Denis Schapiro, Clarence Yapp, Jeremy Muhlich, Artem Sokolov, et al. Highly multiplexed immunofluorescence images and single-cell data of immune markers in tonsil and lung cancer. Scientific Data, 6(1):1–10, 2019.

[28] Daniel Swafford and Santhakumar Manicassamy. Wnt signaling in dendritic cells: its role in regulation of immunity and tolerance. Discovery medicine, 19(105):303, 2015.

[29] Long Long, Xue Zhang, Fuchun Chen, Qi Pan, Pronnaphat Phiphatwatchara, Yuyang Zeng, and Honglei Chen. The promising immune checkpoint lag-3: from tumor microenvironment to cancer immunotherapy. Genes & cancer, 9(5-6):176, 2018.

[30] Lawrence P Andrews, Ariel E Marciscano, Charles G Drake, and Dario AA Vignali. Lag 3 (cd 223) as a cancer immunotherapy target. Immunological reviews, 276(1):80–96, 2017.

[31] Cinzia Solinas, Chunyan Gu-Trantien, and Karen Willard-Gallo. The rationale behind targeting the icos-icos ligand costimulatory pathway in cancer immunotherapy. ESMO open, 5(1):e000544, 2020.

[32] Fei Tu, Ya-Hui Ding, Xi-Hui Ying, Fa-Zong Wu, Xin-Mu Zhou, Deng-Ke Zhang, Hai Zou, and Jian-Song Ji. Regulatory t cells, especially icos+ foxp3+ regulatory t cells, are increased in the hepatocellular carcinoma microenvironment and predict reduced survival. Scientific Reports, 6:35056, 10 2016. doi: 10.1038/srep35056.

[33] Leeat Keren, Marc Bosse, Diana Marquez, Roshan Angoshtari, Samir Jain, Sushama Varma, Soo-Ryum Yang, Allison Kurian, David Van Valen, Robert West, et al. A structured tumorimmune microenvironment in triple negative breast cancer revealed by multiplexed ion beam imaging. Cell, 174(6):1373–1387, 2018.

[34] Alec Radford, Luke Metz, and Soumith Chintala. Unsupervised Representation Learning with Deep Convolutional Generative Adversarial Networks. arXiv e-prints, art. 1511.06434, Nov 2015.

[35] Zhou Wang, Alan C Bovik, Hamid R Sheikh, Eero P Simoncelli, et al. Image quality assessment: from error visibility to structural similarity. IEEE transactions on image processing, 13 (4):600–612, 2004.

[36] Yossi Rubner, Carlo Tomasi, and Leonidas J Guibas. The earth mover’s distance as a metric for image retrieval. International journal of computer vision, 40(2):99–121, 2000.

[37] Kaiming He, Xiangyu Zhang, Shaoqing Ren, and Jian Sun. Deep residual learning for image recognition. In Proceedings of the IEEE conference on computer vision and pattern recognition, pages 770–778, 2016.

[38] Takeru Miyato, Toshiki Kataoka, Masanori Koyama, and Yuichi Yoshida. Spectral normalization for generative adversarial networks. CoRR, abs/1802.05957, 2018. URL http://arxiv.org/abs/1802.05957.

[39] Xudong Mao, Qing Li, Haoran Xie, Raymond YK Lau, Zhen Wang, and Stephen Paul Smolley. Least squares generative adversarial networks. In Proceedings of the IEEE International Conference on Computer Vision, pages 2794–2802, 2017.

[40] S. C. Angelo, M.and Bendall, R. Finck, M. B. Hale, C. Hitzman, A. D. Borowsky, R. M. Levenson, J. B. Lowe, S. D. Liu, S. Zhao, Y. Natkunam, and G. P. Nolan. Multiplexed ion beam imaging of human breast tumors. Nature Medicine, 20:436–442, 2014.

[41] Satelli Arun and Li Shulin. Vimentin as a potential molecular target in cancer therapy or vimentin, an overview and its potential as a molecular target for cancer therapy. Cell Mol Life Sci, 68(18):3033–3046, 2011.

[42] Martha E Kidd, Dale K Shumaker, and Karen M Ridge. The role of vimentin intermediate filaments in the progression of lung cancer. American journal of respiratory cell and molecular biology, 50(1):1–6, 2014.

